# Global Metabolome Analysis of *Dunaliella tertiolecta, Phaeobacter italicus R11* Co-cultures using Thermal Desoprtion - Comprehensive Two-dimensional Gas Chromatography - Time-of-Flight Mass Spectrometry (TD-GC×GC-TOFMS)

**DOI:** 10.1101/2021.09.27.461748

**Authors:** Michael D. Sorochan Armstrong, René Oscar Arredondo Campos, Catherine C. Bannon, A. Paulina de la Mata, Rebecca J. Case, James J. Harynuk

**Affiliations:** Department of Chemistry, University of Alberta, 11227 Saskatchewan Dr NW, Edmonton, T6G 2G2, Alberta, Canada; Department of Human Ecology, University of Alberta, 302 Human Ecology Building, Edmonton, T6G 2N1, Alberta, Canada; Department of Biology, Dalhousie University, 1355 Oxford Street, Halifax, B3H 4R2, Nova Scotia, Canada; Singapore Centre for Environmental Life Sciences Engineering (SCELSE) and School of Biological Sciences, Nanyang Technological University, 60 Nanyang Drive, SBS-01N-27, Singapore, 637551

**Keywords:** Metabolomics, Microalgae, Biofuels, Machine Learning, Chemometrics, GC×GC

## Abstract

*Dunaliella tertiolecta* is a marine microalgae that has been studied extensively as a potential carbon-neutral biofuel source [1]. Microalgae oil contains high quantities of energy-rich fatty acids and lipids, but is not yet commercially viable as an alternative fuel. Carefully optimised growth conditions, and more recently, algal-bacterial co-cultures have been explored as a way of improving the yield of *Dunaliella tertiolecta* microalgae oils. The relationship between the host microalgae and bacterial co-cultures is currently poorly understood. Here, a complete workflow is proposed to analyse the global metabolomic profile of co-cultured *Dunaliella tertiolectra* and *Phaeobacter italicus R11*, which will enable researchers to explore the chemical nature of this relationship in more detail.

**Graphical Abstract:** 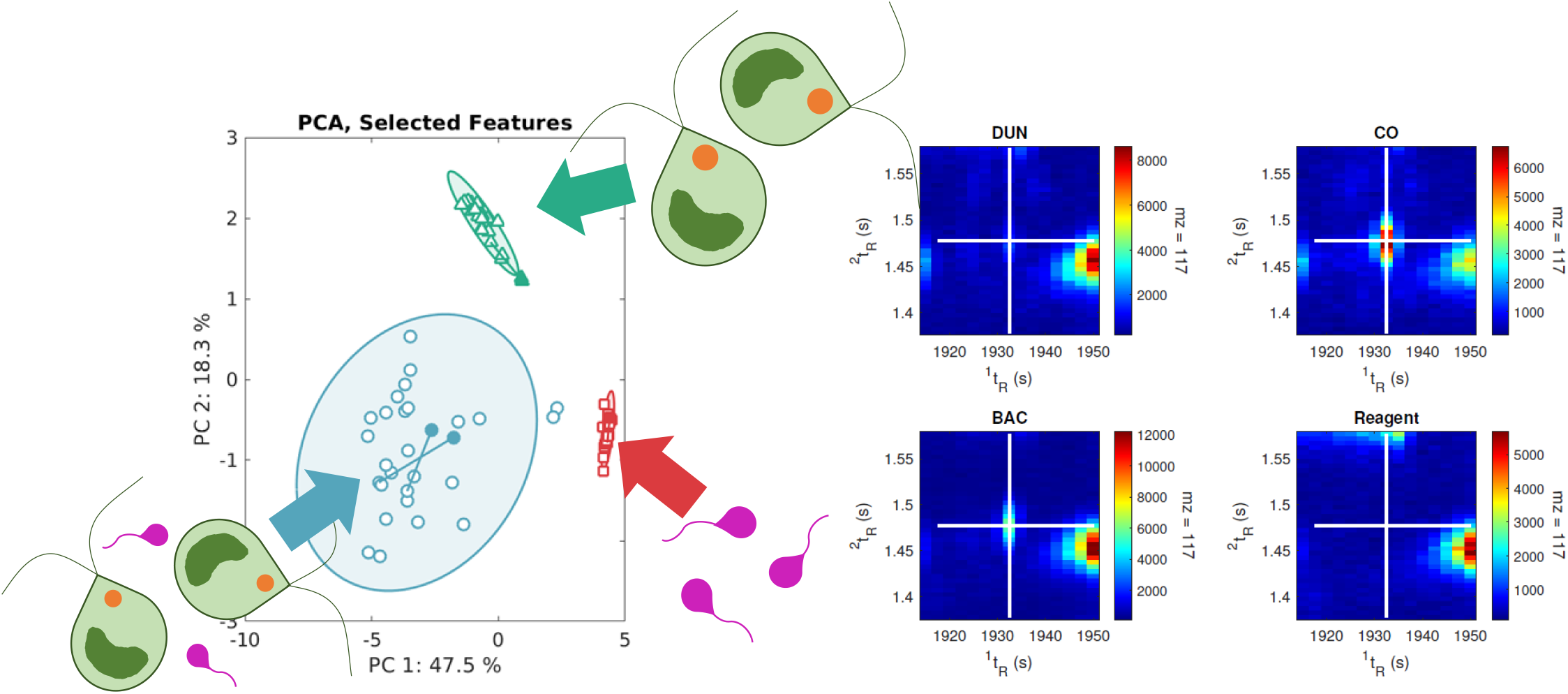

**Highlights:**

- A method for direct microalgae sample introduction is proposed.
- Advanced chemometric tools can extract, useful and discriminating metabolomic features even from very noisy complex datasets.

## 1. Introduction

Microalgae oils represent a source of energy-rich fatty acids and lipids that have long been considered as a carbon-neutral source of biologically-derived fuels [1] [2]. Limiting the commercial applicability of microalgae oil are the long times required for algal growth and the limited yields of oil. In addition to optimisation of growth conditions, bacterial co-cultures have been explored as a way of improving yields of microalgae oils [3]. These co-cultures have improved the rate of growth for microalgae colonies via directly observable mechanisms such as the synergistic exchange of oxygen and carbon dioxide for aerobic bacteria[4], or the release of extracellular compounds into the growth media [5]. For many co-cultures, although a change in the microalgal growth is apparent, there is no clear mechanism that can be inferred through analysis of the growth media alone. In these cases, it may be useful to examine the biomass directly.

Metabolomics has been used to great effect to correlate differences in small-molecule metabolite expression with macroscopically observable phenomena. Algae metabolomics in particular has seen some interest [6] [7], and has been used to gain insight into chemical responses of microalgae to changes in their environment. The relative expression of these chemicals can be used to deduce the ecology of the bacterial-algal relationships as either mutualistic [8] [9] or antagonistic [10], and such insight may be used to further improve upon the microalgae oil yield.

Comprehensive two-dimensional gas chromatography - time-of-flight mass spectrometry (GC×GC-TOFMS) is a powerful analytical tool for the non-target examination of the chemical diversity of samples of volatile and semi-volatile organic compounds owing to its improved resolution, sensitivity, and identification capabilities over traditional one-dimensional gas chromatography - mass spectrometry (GC-MS). For non-volatile species and those which chromatograph poorly, such as lipids, fatty acids, and amino acids, extra derivatisation steps are required. These steps hydrolyze bonds in large molecules (e.g. triacyl glycerides) and substitute groups such as methyl-, ethyl-, or trimethylsilyl-for labile protons, enabling subsequent analysis in the gas-phase. Techniques specific to a particular class of chemical compound, usually fatty acids, are common in algae metabolomics since only a limited number of chemical classes dominate the composition of microalgae oil. Analysis of Fatty Acid Methyl Esters (FAMEs) is a common way of determining fatty acid expression by GC-MS, for example. Targeted analysis of a limited number of chemical species often fails to explain the observed phenomena. Global metabolomic profiling aims to encompass the broadest possible scope of all small-molecule metabolites, for the similar aim of correlating changes in abundances of some small number of metabolites with the different classifications of populations comprising the study data set (e.g.: healthy/diseased or different cultures of microalgae).

GC×GC-TOFMS often reveals several thousands of unique chemicals in a metabolomics study. For a properly optimised analysis, most of these chemicals originate from the biomass itself. However, contamination at some stage of the sample preparation is inevitable; especially for samples requiring derivatisation, as the reagents involved are particularly difficult to purify. Even for chemical features that are biological in origin, typically only a small subset of these features contain useful or discriminating information. As such, for a limited number of samples, a feature selection step is necessary to create informative models that are easy to interpret.

In this submission, the authors present a workflow for preparing microalgae samples for global metabolomic analyses that includes a novel technique for introducing the derivatised sample matrix into the gas chromatograph. Using thermal desoprtion (TD), a concentrated sample of microalgae extract can be introduced directly into the instrument without a prior filtration step. The injected sample is deposited into a small insert in the thermal desoprtion unit, and heated to transfer the volatile and semi-volatile components to a cryogenically cooled inlet. All low-volatility components that would otherwise be deposited into the inlet itself remain within the insert inside of the TD unit. Subsequent pyrolysis of heavy biomass residues between sample injections is avoided in this way, and the analyses are relatively free of interference and contamination. 70 samples of *Dunaliella tertiolecta, Phaeobacter italicus R11* [11] [12], and co-cultures of the two species were cultured and filtered for analysis. A useful sample normalisation and feature selection routine is demonstrated on this dataset, and the authors present a short list of candidate metabolites that may be biologically interesting for future study.

## 2. Materials and Methods

### 2.1. Preparation of Dunaliella tertiolecta, Phaeobacter italicus R11, and co-cultured samples

#### 2.1.1. Growth and maintenance of algal and bacterial strains

The *Dunaliella tertiolecta* CCMP 1320 strain was obtained from the Provasoli-Guillard National Centre for Marine Algae and Microbiota (NCMA). The chlorophyte was maintained in L1-Si media made with artificial seawater (35 g/L of Instant Ocean, Blacksburg, VA, USA), at 18 °C with a diurnal incubator cycle (12:12 hour dark-light cycle). Samples from the cultures of microalgae were examined microscopically to rule out bacterial contamination before experimental use. These samples were also inoculated onto marine agar plates, and incubated at 28°C for 3 days to identify any colony forming units (CFUs). (18.7 g of Difco Marine Broth 2216 with 9 g NaCl and 15 g Difco agar in 1 L). Experimental use of the algal cultures proceeded once a cell concentration of 10^4^ cells/mL was reached.

Samples of *Phaeobacter italicus R11* were acquired from Botany Bay, Australia [11]. The bacterial cultures were maintained at 28 °C on the aforementioned marine agar plates, then transferred to 5 mL 50% dilute marine broth media (2216 Marine Broth, Difco) where it was grown until reaching a stationary phase for 24 hours before the experiments. Cell concentration for stationary phase *Phaeobacter italicus R11* was similar to the cell concentration of the algae cultures, at 10^4^ cells/mL.

#### 2.1.2. Preparation of samples

Algal-bacterial co-cultivation experiments were performed as described by Bramucci et al. [13] in 12 well plates (Standard TC Growth Surface, Bacto (Oakville, Canada)). Briefly, stationary phase bacterial colonies of *Phaeobacter italicus R11* R11 were washed twice by centrifugation and re-suspended in L1-Si medium before dilution to the target cell concentration 10^4^ colony-forming units (CFU) /mL. For co-culture samples, *D. tertiolecta* and *Phaeobacter italicus R11* were mixed in a 1:1 ratio by volume at equivalent cell concentrations in L1-Si medium made with artificial seawater. Mono-culture controls of both *D. teriolecta* and *Phaeobacter italicus R11* R11 were inoculated in 1:1 (v:v) ratio with L1-Si made with artificial seawater. Mono- and co-cultures were aliquoted in 6 mL volumes into 12-well plates and grown in a diurnal incubator with a 12 hr dark-light cycle at 18°. During the mid-point of each dark cycle, 20 µL of each sample were plated onto a 1.5% agar, 1/2 marine broth plates and incubated for 24 h at 28°C to confirm the absence of bacterial in monoculture samples, and enumerate the bacteria in co-culture samples. All samples were cultured until the stationary phase, after 18 days.

#### 2.1.3. Collection of culture samples

On day 18 of each samples’ incubation, each sample was collected by vacuum filtration onto pre-weighed glass fibre filters (0.22 *µ*m). Each sample was rinsed three times with 1 M PBS buffer to wash away the growth media. Filters were placed into clean, glass vials covered with Kimwipes and stored at -80 ° before being lyophilised for 24 hours. After the drying step, the filters were weighed once again to record the biomass. The biomass was recorded to (± 0.001 g), which was insufficient to accurately determine the mass of either bacterial culture samples, or the algae and culture samples. The bacterial samples were recorded at 0.000 ± 0.002 g. Co-culture, and *D. tertiolecta* samples presented an average biomass of 0.005 g ± 0.002 g and 0.005 g ± 0.002 g respectively.

20 samples of *D. tertiolecta* + 1 replicate, quality control samples, 23 samples of *Phaeobacter italicus R11* + 1 replicate, quality control sample, and 27 co-cultured samples + 3 replicate quality control samples were prepared and analysed for the experiment. One quality control sample, part of the co-cultured samples, indicated poor agreement with its corresponding sample, and indicated a change in the tightly-controlled analytical conditions of the instrument. This sample was excluded from the final dataset, bringing the total number of samples (including quality controls) to 74. Careful inspection of data for samples before and after this particular sample indicated that this one anomalous result was the result of an isolated incident that did not affect other samples.

### 2.2. Sample preparation and derivatisation

Liquid measurements were performed using either a 100-*µ*L or 1000-*µ*L Microman positive displacement pipette and disposable pipette tips. For measurements of pure reagent, a reusable positive displacement syringe system with a digital readout was used (SGE eVol^*TdM*^ handheld automated analytical syringes).

The sample preparation procedure is divided into two main steps. During the first step, an extraction protocol based on the widely-known Bligh and Dyer [14] [15] method for the analysis of fatty acids was used to extract and separate the macro-molecular plant material from the plant metabolites. Use of the chloroform extract appeared to be a good choice of extraction solvent and offered decent coverage of a wide variety of different chemical classes. During the second step, trimethylsilyl derivatives of the extracts were generated based on the protocol of Chan et al. [16].

The glass fibre filter papers (GFFPs) used to collect each sample were submerged in 7-mL volumes of HPLC-grade methanol (>99.9%, Millipore-Sigma Canada) in 20-mL scintillation vials (Chromatographic Specialties Inc., Oakville, ON, Canada) using a clean spoonula. To each vial, 7 mL of HPLC-grade chloroform (>99.8%, Millipore-Sigma Canada, Oakville, ON, Canada) were added, and sonication proceeded for an additional hour. 3.5 mL of 18.2 MΩ deionised water (Elga PURELAB flex 3 system, VWR International, Edmonton, AB, Canada) was then added to effect a separation into a polar methanol/water layer and a chloroform layer, which was allowed to rest overnight at 3 °C prior to extraction. 1.8 mL of the bottom (chloroform) layer was extracted and transferred into 2-mL glass GC vials. For replicate measurements, an additional 1.8 mL was transferred to an additional vial. The extracts were blown down with nitrogen at 40 °C until there was no visible chloroform remaining, approximately 2 hours. 100 *µ*L of toluene was added to the dry residue, which was then vortexed briefly before being evaporated under nitrogen at 40 °C once again. The extra evaporation step using toluene ensures there no remnants of moisture remained in the vials as even traces of moisture will interfere with the subsequent derivatisation steps. 50 *µ*L of methoxyamine (Millipore Sigma, Oakville, ON, Canada) in HPLC-grade pyridine (Millipore Sigma, Oakville, ON, Canada) was added via a digital positive displacement syringe, and the solution was incubated at 60 °C for 2 hours. Following this, using another positive displacement syringe, 100 *µ*L of N-Methyl-N-(trimethylsilyl)trifluoroacetamide + 1% trimethylchlorosilane (MSTFA + 1% TMCS, Fisher Scientific, Ottawa, ON, Canada) was added and vials were incubated at 60 °C for an additional hour. 100 *µ*L of the resultant solution was transferred to 1.8 mL vials with fused glass 300-*µ*L inserts. These vials were stored at 3 °C for up to 48 hours prior to analysis.

### 2.3. Sample introduction and operating conditions

#### 2.3.1. Thermal desorption, sample introduction

The chloroform extracts were not centrifuged to remove non-volatile components from the extract solution. Being a relatively non-polar solvent, exposure of chloroform to plastic centrifuge tubes would present a significant risk of leeching contaminants from the vials into the samples. To avoid this, the authors opted to inject the unfiltered, derivatised extract directly into TDU insert tubes, with subsequent evaporation of (semi-)volatile components directly from the insert tube, leaving the non-volatile components in the insert. Inserts were replaced for every analysis.

A Gerstel MPS autosampler and sample preparation robot equipped with a 10-*µ*L Gerstel TriStar Liquid Syringe, a Thermal Desoprtion Unit (TDU 2), and a programmed temperature vaporization inlet (CIS4; Gerstel US - 701 Digital Drive, Suite K, Linthicum, MD 21090). The MPS system was programmed in Automated TDU-Liner Exchange (ATEX) mode, where 9-*µ*L aliquots of sample were injected into clean TDU tubes containing a disposable microvial insert. TDU tubes (straight tubes with notch), were rinsed with high-purity (99.9%) toluene (Millipore Sigma Canada, Oakville, Ontario) and baked in an oven at 400 °C for 1 hour between runs, with new microvial inserts baked inside of the TDU tubes. Both the microvial inserts and TDU tubes were allowed to cool inside of the oven before being fitted with transport adapters for liquid injections. The transport adapter seals the microvial inside of the TDU tube, maintaining cleanliness. A Teflon-coated septum in the adapter maintains carrier gas pressure before and after the liquid injections. The liquid syringe was washed 6 times with each of 10 *µ*L of 1:1 (v/v) acetone:hexane and HPLC grade methanol (Fisher Scientific Co, Edmonton Alberta) both before and after the liquid injection.

The TDU was operated in solvent vent mode, followed by a splitless injection from the TDU to the inlet, and then splitless injection from the inlet to the GC×GC-TOFMS. During the solvent vent step, the TDU was kept at an initial temperature of 128 °C, and the TDU split vent was open for 5 min to vent the pyridine and MSTFA solvent mixture. The TDU was fed with a constant flow supply of ultra-high purity helium carrier gas (Linde Canada (formerly Praxair), Edmonton Alberta, Canada) at 50 mL/min with the permanent split vent from the TDU set to 2.5 mL/min. The remaining flow during the solvent vent step exited the system via the split line from the CIS. Following solvent venting, the temperature of the TDU was raised to 280 °C and injected via a splitless injection into the CIS at a flow rate of 50 mL/min for an additional 5 min. During this step, the CIS was maintained at a temperature of 30 °C with the split valve open, until the splitless injection from the TDU was completed. 30 °C was the lowest practical temperature that could be reliably maintained with the cryogenic cooling system during the run. In the next step, the CIS ramped to 300 °C and injected with the split vent closed into the GC×GC-TOFMS. During this time, the TDU ramped to 300 °C to clean the system, with the TDU split vent open. The CIS operated in splitless mode for 150 s at 300 °C, until the split vent was opened, at a total gas flow rate of 252 mL/min for a constant flow rate of 2 mL/min delivered to the head of the column. The inlet liner was baffled (Gerstel, US) to trap volatile components, while allowing for an effective purging step between each run, and system cleanliness was monitored via instrument blanks (one fast blank after every sample, and one instrument blank under identical operating conditions to those used for the samples twice per batch of 14 samples).

#### 2.3.2. GC×GC-TOFMS method

The samples were separated on a LECO Pegasus 4D system (LECO, St. Joseph, MI, USA) outfitted with a quad-jet dual-stage cryogenic modulator. The column set featured a 60 m × 0.25 mm internal diameter; 0.25 *µ*m film thickness Rxi-5SilMS in the first dimension, and a 1.4 m × 0.25 mm internal diameter; 0.25 *µ*m film thickness Rtx-200MS second dimension column (Chromatographic Specialities, Brockville, ON, Canada). The initial oven temperature was set to 80 °C, held for 4 min, and ramped at 3.5 °C/min to a maximum oven temperature of 315 °C with a 10 min final hold. The temperature program and flow rate of the method were directed by considerations for speed optimised flow (SOF) [17] and optimal heating rate (OHR) [18] derived from the column geometry and dead-time, respectively. The secondary oven offset was set at +10 °C relative to the primary oven temperature and the modulator temperature offset was set at +15 °C relative to the secondary oven temperature. The modulation period (*P*_*M*_) was 2.50 s, with a hot pulse time of 0.60 s and a cool time of 0.65 s between stages. The mass spectrometer collected spectra at 200 Hz, from 40 to 800 (m/z). The election impact ionisation energy was -70 eV. The ion source temperature was 200 °C, with a transfer line temperature of 300 °C. An acquisition delay of 650 s was used to ensure residual solvent did not damage the filament.

#### 2.3.3. GC×GC-TOFMS data pre-processing

The data were pre-processed using LECO ChromaTOF^®^ version 4.72. Baseline offset for peak detection was set as a factor of 1.2 above the estimated noise level. Anticipated peak widths were determined through a survey of 10 different peaks, both large and small. The authors opted to direct the pre-processing parameter optimisation towards detecting smaller peaks, so the peak size parameters were as follows: 10 s for first-dimension peak width, and 0.1 s for second-dimension peak widths. The second-dimension sub-peaks were combined if their deconvolved mass spectra met a match factor of 650, and sub-peaks were only integrated if their signal-to-noise ratio (SNR) was greater than a value of 6.

Peaks were integrated in the final peak table, if the SNR for the base peak was greater than a value of 15, with 5 or more apexing masses. Peaks across multiple samples were aligned if they were within two modulation periods of each other in the first-dimension, or within 0.2 s of each other in the second-dimension. Quality control samples were included in the alignment procedure, bringing the total number of samples to 75 (including one sample that was later discarded). Before being included in the final table, an analyte must have populated at least 33% of the samples (25 samples), otherwise that analyte was discarded. This parameter was selected to minimise the very common phenomenon of peak dropout in processed GC×GC-TOFMSpeak tables, and was not based on the population of unique classes of sample. Since the number of variables for typical GC×GC-TOFMSexperiments is much higher than the number of samples, spurious class separations for targeted classification are very common for poorly optimised peak tables. However considering the class makeup of the dataset, chemical components unique to *D. tertiolecta* and co-cultured samples, *Phaeobacter italicus R11* and co-culture samples, as well as the co-culture class itself, are not disallowed from the final peak table under these conditions.

Peak searching for analytes not recognised in the sample-wise pre-processing were searched for down to a SNR ratio of 12.5. Initial library searching for all peaks was performed for all samples using the National Institute of Standards and Technology (NIST) mass spectral database (2017).

### 2.4. Data analysis

The final peak table was exported from ChromaTOF^®^, and into MATLAB^®^ 2020b. Some functionalities of PLS Toolbox (R8.5.2; Eigenvector Research Inc.) were used for the receiver-operator characteristics (ROC), as was the MATLAB^®^ Statistics and Machine Learning Toolbox. All data used in this work are available online at https://doi.org/10.20383/102.0510.

#### 2.4.1. Sample normalisation

Samples were normalised according to a class-based total useful peak area (cTUPA) criterion, based on the absence of a reliable biomass measurement, and the need to account for variability in the quantity of sample that eventually reached the instrument. Using this method, peak areas were divided by the total peak area of analytes detected in each sample of a given class. This allowed for a reasonable, and simple method for comparing highly dissimilar samples (i.e. comparing a small bacterial biomass against a much larger algal biomass) and extracting a useful discriminating variable subset that best describes their differences. A similar approach, agnostic to class labels, was reported for a study conducted on GC×GC-TOFMS using human urine [19].

#### 2.4.2. Feature selection, cross-validation

The normalised data were analysed using feature selection by cluster resolution (FS-CR), a hybrid wrapper/threshold method for selecting features based on favourable projections within a principal component subspace. This technique has shown to be effective on a number of GC×GC-TOFMS datasets, and includes a robust cross-validation routine that trains, optimises, and validates the model during each iteration. To demonstrate the effectiveness of this tool, a series of ROC curves were generated by reshuffling the data for each iteration, utilising a variable subset that survived a certain ratio of previous combination of the data. A PLS-DA model with 2 latent variables was trained using half of the reshuffled samples, and validated using the other half of the reshuffled samples. Further details are presented in the Results section.

The FS-CR algorithm was operated for projection into two-dimensional principal component subspaces, using autoscaled data (i.e. data that was mean-centred and scaled via each variable’s standard deviation). The most recent version of the algorithm was selected, utilising a numerical determination of the cluster resolution metric[20], which allowed for 200 iterations of the algorithm to complete in about 30 min. Variables that survived 90% of all iterations were ultimately selected for the final model.

Mass spectra and retention indices for analytes of interest were recovered from the raw data and a library search using the Golm Metabolome Database [21] was performed.

## 3. Results

Each of the resultant chromatograms are rich in chemical information, but at first glance they are visually quite similar. The colour axes of Figures 1-3 are scaled to the same maximum TIC of 7.5 × 10^5^. In spite of the high sample load, instrument blanks were clean between runs, but despite the authors’ best efforts, the reagent blank itself a great deal of interfering chemical information (See: Supporting Information). This is likely unavoidable, as the purity of MSTFA used for derivatisation is generally quite poor (98%), and the pre-concentration step during the sample introduction likely exacerbated this problem.

**Figure 1:**
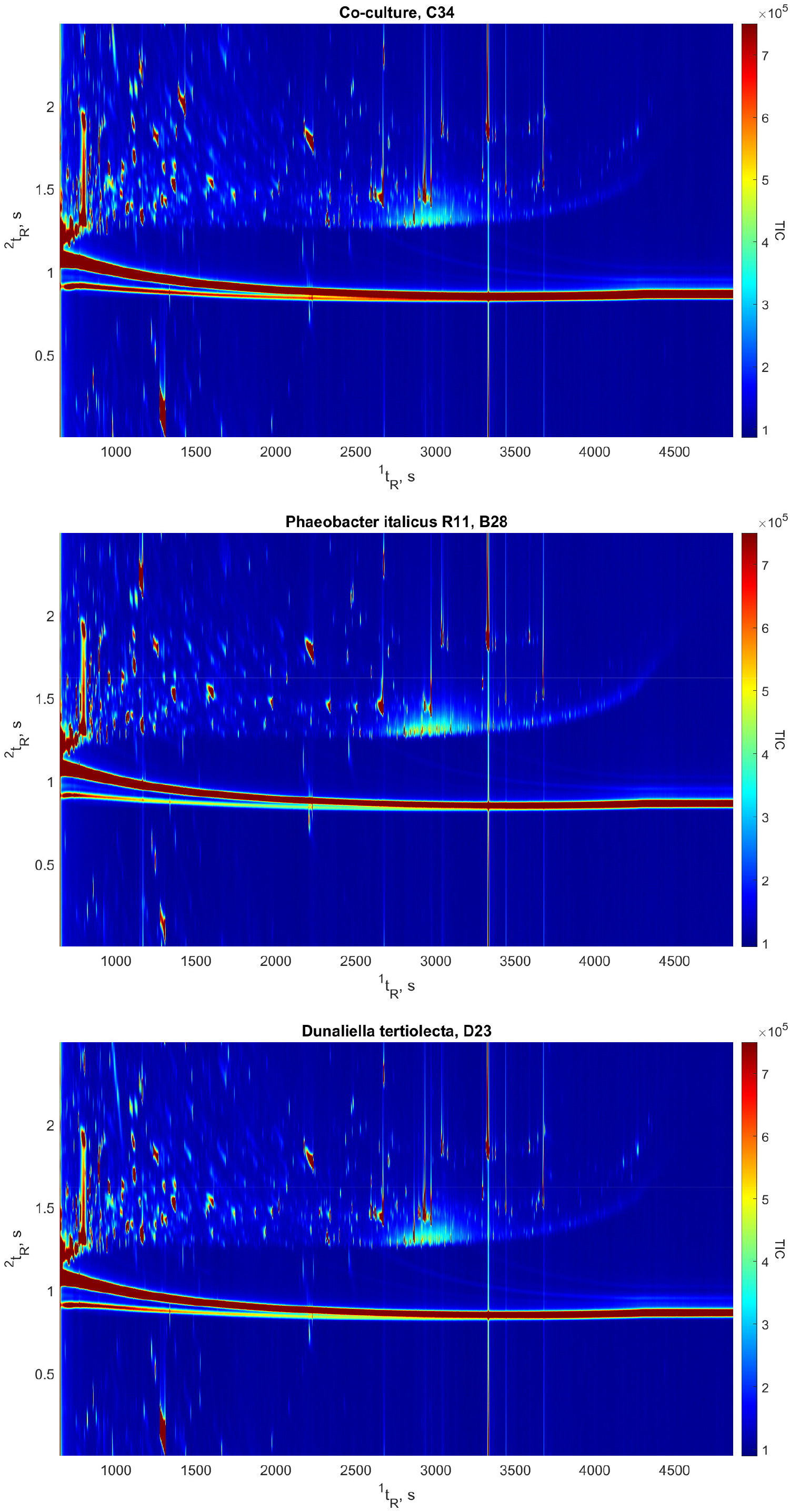
Example Total Ion Current (TIC) chromatograms from each sample class

Prior to analysis, all data was autoscaled. For the ROC curves, validation data was centered and scaled according to the values obtained in the training set, for a more critical assessment of the model performance. The principal component analysis of the raw data revealed some bimodality, likely due to the number of interfering chemicals present in the reagent blank. Following cTUPA, the severity of the bimodality was reduced, and the cluster of bacterial samples was well resolved from the chemically similar co-culture and mono-culture microalgae samples (Figure 2) This aligns well with our initial expectations of the data.

**Figure 2:**
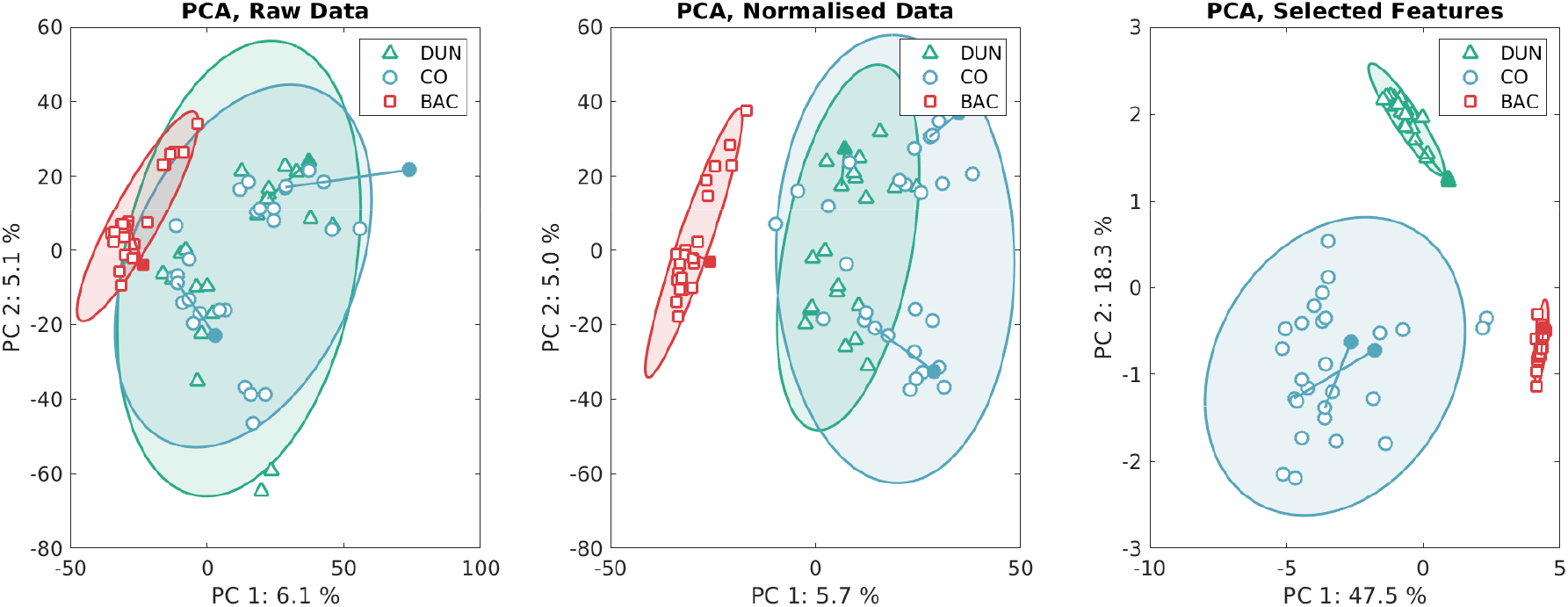
From left to right: results of principal component analysis of the raw data (autoscaled), similarly scaled data normalised to class-specific TUPA, and the normalised, scaled data using the selected features from the FS-CR routine. Quality control samples were not included in the feature selection routine, and are displayed as filled icons connected to their corresponding replicate with a straight line, following projection into the optimised principal component space.

Using the selected features, the samples appear to be normally distributed about the two axes of variance within each cluster. This suggests that the extracted features are robust against interfering chemical information, and that the previously observed bimodality of the data may have been a chemical, or instrumental artefact of the analysis that did not significantly affect projections using the selected features (Figure 2).

In-class variance within the co-cultured microalgae samples appears to be much higher than in the mono-culture samples, according to the projection within the subspace of the selected features. Conversely, in-class variation for the bacterial and mono-culture classes is relatively low, suggesting the extracted metabolites are disregulated within co-cultured samples (Figure 2).

The cross-validated receiver operator characteristics support the utility of this feature selection routine, and the extracted chemical characteristics. Although external validation was not used, regardless of the samples chosen, the calculated area under the curve is very close to ideal for all 200 combinations of the data that were independently selected relative to the feature selection routine (Figure 3).

**Figure 3:**
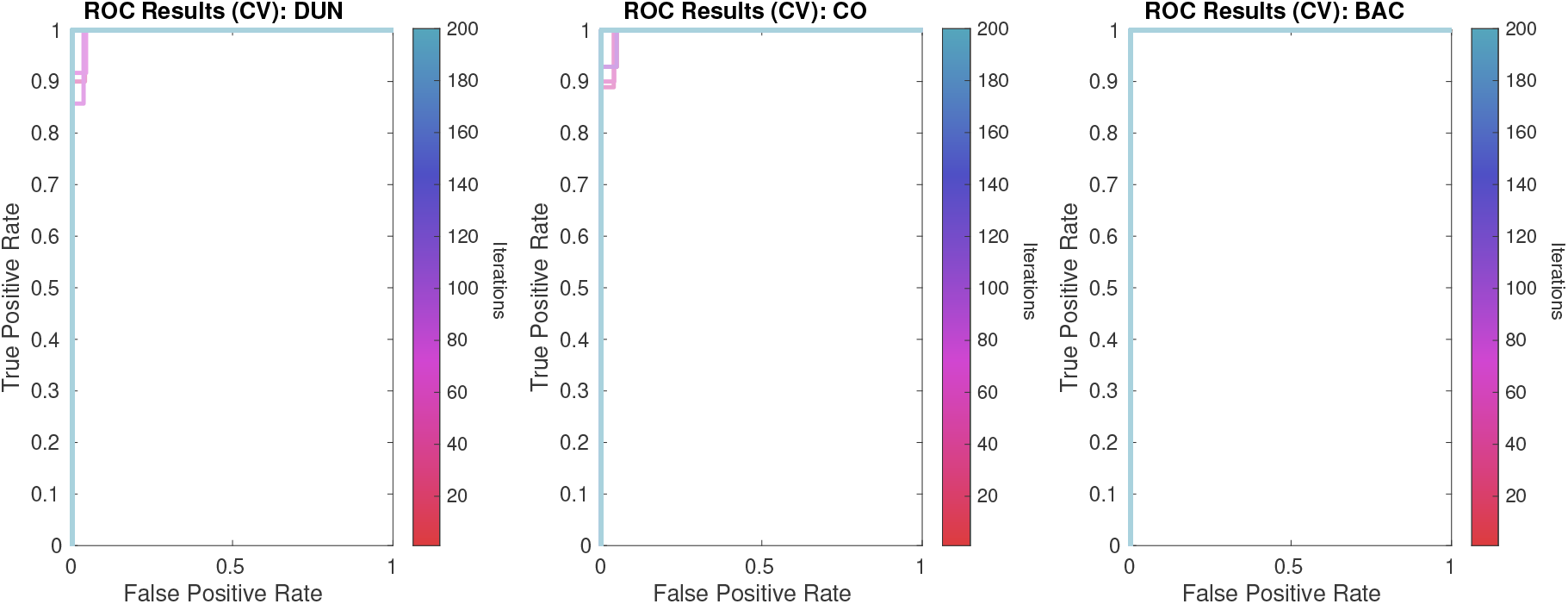
Cross validated receiver operator characteristics suggest that the calculated model is robust, and improves with more iterations. Light blue lines show the results of further iterations, and red lines show the results of fewer iterations. Each line is semi-transparent, but the AUC is close to 1 in all cases. Calculation of the classification scores was based off the named class (Class 1), versus everything else (Class 0).

## 4. Discussion

16 analytes of interest were selected using the feature selection routine. In the supporting information, there is a summary of each extracted analyte with the most relevant information presented. Using each analyte’s quantification ion listed in the output of ChromaTOF^®^, it is appears that the majority of the selected features are not spurious signals, given their roughly Gaussian, uni-modal peak profiles in both dimensions. However, the quality of the deconvolution appears to be poor in some features that returned no probable library hit. This can be observed by the lack of isotopic mass distributions for many prominent peaks in the mass spectra. It’s unlikely that these features can be identified based solely on the extracted mass spectra, although that isn’t to say that the features are not reliable.

Some library hits from the Golm Metabolome Database scored reasonably well on the mass spectral dissimilarity score (1-dot product), but did not account for a number of prominent peaks in the observed mass spectra. It’s possible that for these standards, no good library spectra exist. For Analytes 5083 and 22226, a very prominent peak at m/z = 143 is observed. This is a prominent peak in the mass spectra for alkyl-quinolones [22], a class of antibiotics. Analyte 27584 appears to bear some similarity to an unidentified compound uploaded to the Golm database as part of an earlier study [23].

Representative mass spectra were extracted from individual samples where that compound was identified. A drawback of ChromaTOF^®^ software is that each samples’ features present their own mass spectra, and these mass spectra are associated across multiple samples provided that a certain similarity in mass spectral characteristics and retention time is reached. This means that the mass spectra from each sample may differ somewhat, but the decision was made not to average the mass spectra across multiple samples, as doing so could lower the precision of the extracted mass spectra or bias the match score of the library match.

Further study will is needed to confirm the identities of these analytes, since few were identified at level 2 or higher according to the Metabolomics Standards Initiative [24]. Below is a table summarising the quality of each of the mass spectral hits, illustrating the reasoning behind each hits’ identification level. Compounds that were found in the reagent blank in addition to the samples were assumed to be less reliable features, but may nonetheless still be identified.

**Table 1:**
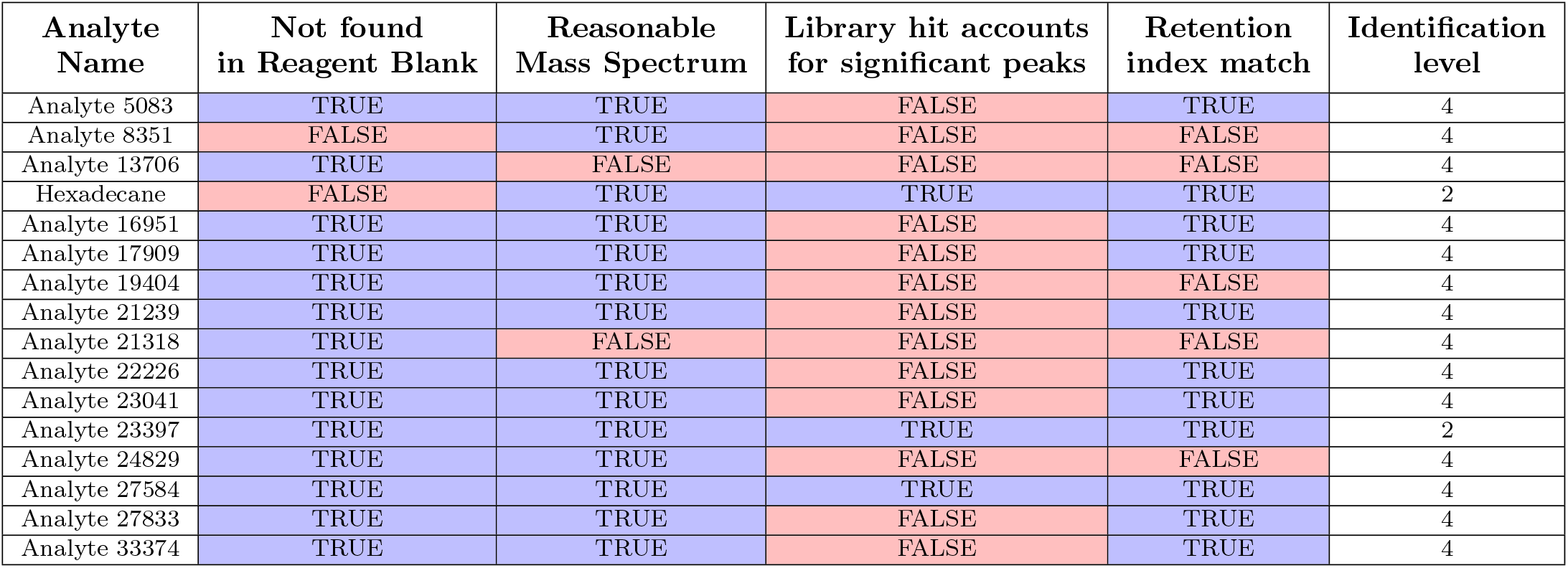
Identification levels for the significant features in the dataset, summarising the factors that went into ascribing MSI Identification levels 2 (punative identification) or 4 (unknown metabolite). Further details are available in the supporting information.

| Analyte Name | Not found in Reagent Blank | Reasonable Mass Spectrum | Library hit accounts for significant peaks | Retention index match | Identification level |
| --- | --- | --- | --- | --- | --- |
| Analyte 5083 | TRUE | TRUE | FALSE | TRUE | 4 |
| Analyte 8351 | FALSE | TRUE | FALSE | FALSE | 4 |
| Analyte 13706 | TRUE | FALSE | FALSE | FALSE | 4 |
| Hexadecane | FALSE | TRUE | TRUE | TRUE | 2 |
| Analyte 16951 | TRUE | TRUE | FALSE | TRUE | 4 |
| Analyte 17909 | TRUE | TRUE | FALSE | TRUE | 4 |
| Analyte 19404 | TRUE | TRUE | FALSE | FALSE | 4 |
| Analyte 21239 | TRUE | TRUE | FALSE | TRUE | 4 |
| Analyte 21318 | TRUE | FALSE | FALSE | FALSE | 4 |
| Analyte 22226 | TRUE | TRUE | FALSE | TRUE | 4 |
| Analyte 23041 | TRUE | TRUE | FALSE | TRUE | 4 |
| Analyte 23397 | TRUE | TRUE | TRUE | TRUE | 2 |
| Analyte 24829 | TRUE | TRUE | FALSE | FALSE | 4 |
| Analyte 27584 | TRUE | TRUE | TRUE | TRUE | 4 |
| Analyte 27833 | TRUE | TRUE | FALSE | TRUE | 4 |
| Analyte 33374 | TRUE | TRUE | FALSE | TRUE | 4 |

Several of listed metabolites appear to be biologically active based on their relative expressions. These metabolites may affect cell-cell signaling, hormone regulation, or be utilised as antibiotics in co-culture samples. These are of particular interest, since bioactive compounds are known to play a role in bacterial-algal interactions. Of particular interest is the homoserine lactone (HSL) which is produced by *Dunaliella tertiolecta*, however it is absent from *Phaeobacter italicus R11* B28 and the co-culture sample C34 (Supporting Information). HSLs are only known to be produced by the bacterial group Proteobacteria [25] and so further structural elucidation is necessary to assert whether or not HSL has actually been produced by a eukaryote. Previously, Schaffer et al have identified a plant metabolite, coumaric acid, that can replace the HSL tail to form a hybrid material-host signal [26], and so an analogous system with the HSL ring structure being host derived warrants further investigation. Alternatively, the Dunaliella teriolecta metabolite could be an HSL antagonist as *Phaeobacter italicus R11* is known to produce HSLs which are antagonised by the algal metabolites, furanores [11] [27] [28].

## 5. Conclusion

Despite considerable interference from chemical and pre-processing artefacts, using advanced instrumentation and data analysis methods it is possible to gain unprecedented insight into the metabolome of commercially interesting microalgae samples. A complete workflow for profiling the global metabolome of microalgae samples has been proposed, that may guide selection of bacterial inoculations for microalgae cultures to improve the yield of cultivated, carbon-neutral biofuels in the future. This study has also demonstrated the feasibility of using thermal desorption as a sample introduction technique that can allow larger-than-normal aliquots of sample to be introduced with effective pre-concentration and clean-up of dirty samples, without the need for a centrifugation step. Additionally, the utility of cTUPA as a normalisation strategy for highly dissimilar sample classes was demonstrated.

Further work is needed to interpret, and test resultant theories of the relationship between *D. tertiolecta* and *Phaeobacter italicus R11* cultures. Doing so may enable cultivation techniques that exploit this relationship for dividends in sustainable fuel development. Presented here is a complete workflow, with some preliminary results, that can serve as a benchmark for future experiments.

## 6. Acknowledgments

The authors wish to acknowledge the support of the Natural Sciences and Engineering Research Council of Canada (NSERC) and the support given to The Metabolomics Innovation Centre (TMIC) through grants from Genome Alberta, Genome Canada, and The Canada Foundation for Innovation. This work was also supported by A*STAR SFS IAF-PP grant (A20H7a0152) awarded to Rebecca Case.

