## Supporting Information for "Global Metabolome Analysis of *Dunaliella tertiolecta, Phaeobacter italicus R11* Co-cultures using Thermal Desoprtion - Comprehensive Two-dimensional Gas Chromatography - Time-of-Flight Mass Spectrometry (TD-GC×GC-TOFMS)"

September 27, 2021

#### Contents

**1 Example Instrument Blank, Reagent Blank** **2**

**2 Overview of Extracted Features** **3**

### 1 Example Instrument Blank, Reagent Blank

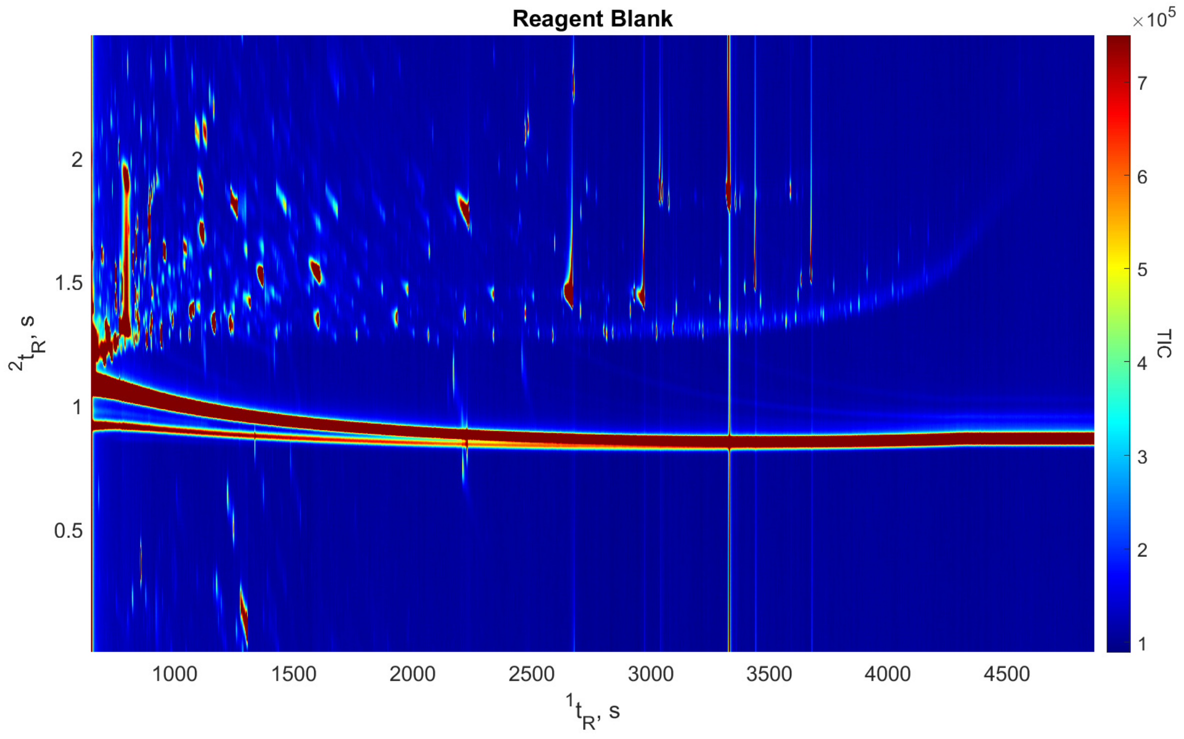


Figure 1: Example reagent blank, demonstrating the low purity of derivatisation reagents. Subsequent analysis will show that despite the high number of interfering chemical components, the extracted features were generally not found in the reagent blanks.

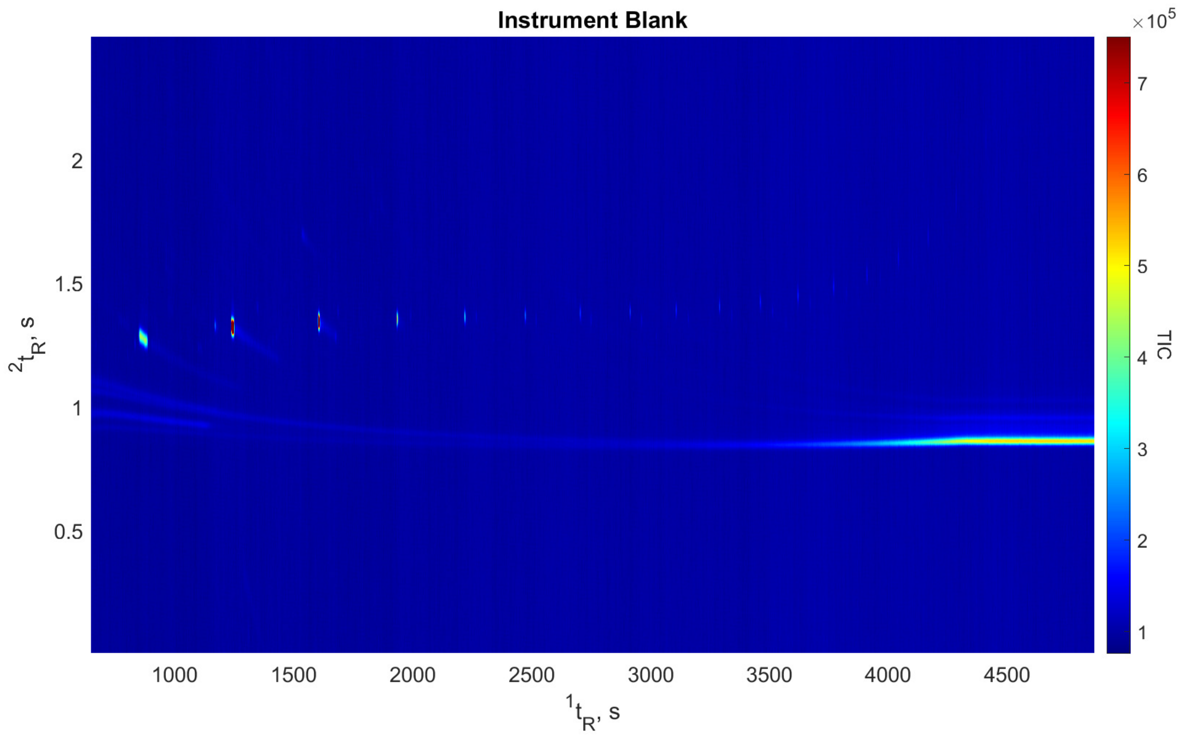


Figure 2: Example instrument blank, demonstrating that the instrument was largely free of contamination between runs, despite a heavy sample load. The observed peaks are cyclic siloxanes, a byproduct of column degredation. These peaks were not identified as significant in the analysis of the data.

### 2 Overview of Extracted Features

Below are the summary analyses of the features determined to be most significant according to the FS-CR routine. For each peak, the most relevant details about the features are presented. The chromatograms display the peak using the quantitative ion identified by ChromaTOF, with the nominal retention times indicated by the white cross. A representative sample for each class is displayed, including a reagent blank. A plot comparing the mass spectra, in addition to a box plot summarising the significance of each feature per class is also presented.

Linear alkanes between $n = 12$ and $n = 32$ were run at the start of the analysis, and retention indices were calculated for all analytes that eluted within this window - otherwise the retention index is listed as undefined.

Poor library matches within the Golm Metabolome Database do not reflect the quality of the selected metabolites. The library may not contain an appropriate standard, or the library mass spectra may have been collected on a different type of mass spectrometer. Definitive identification of these analytes remains an open research avenue for future work.

| Peak Information | Top Golm Metabolome Database Search Result |
|---|---|
| ${}^{1}t_R$(s): 967.5 | |
| ${}^{2}t_R$(s): 2.070 | |
| Quantification Ion (m/z): 143 | 1-dotprod (Spectral Dissimilarity): 0.1923 |
| Analyte Name (ChromaTOF): Analyte5083 | Analyte Name (Library): Alanine, beta- (1TMS) |
| Retention Index (Observed): Undefined | Retention Index Difference (Library): Undefined |

**Link:** http://gmd.mpimp-golm.mpg.de/Spectrums/784e1232-c517-423f-98f9-ae3eb5351dac.aspx
**Contributor:** Jäger C, Schomburg D Department of Bioinformatics and Biochemistry, Technische Universität Carolo-Wilhelmina Braunschweig, Langer Kamp 19B, D-38106 Braunschweig, Germany

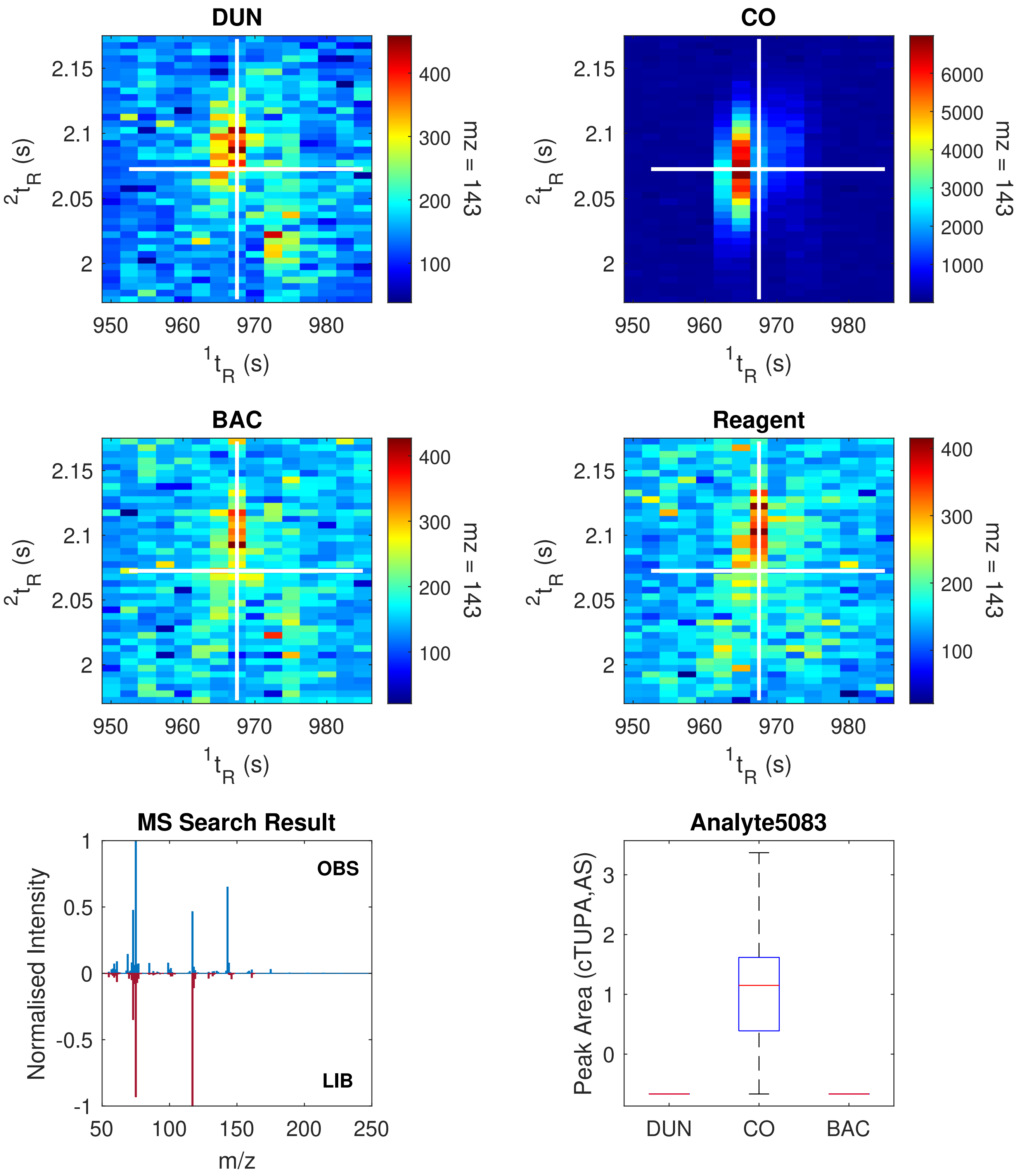

| Peak Information | Top Golm Metabolome Database Search Result |
|---|---|
| $^{1}t_R$(s): 1212.5 | |
| $^{2}t_R$(s): 1.695 | |
| Quantification Ion (m/z): 203 | 1-dotprod (Spectral Dissimilarity): Unknown |
| Analyte Name (ChromaTOF): Analyte8351 | Analyte Name (Library): Unknown |
| Retention Index (Observed): 1284.99 | Retention Index Difference (Library): Unknown |

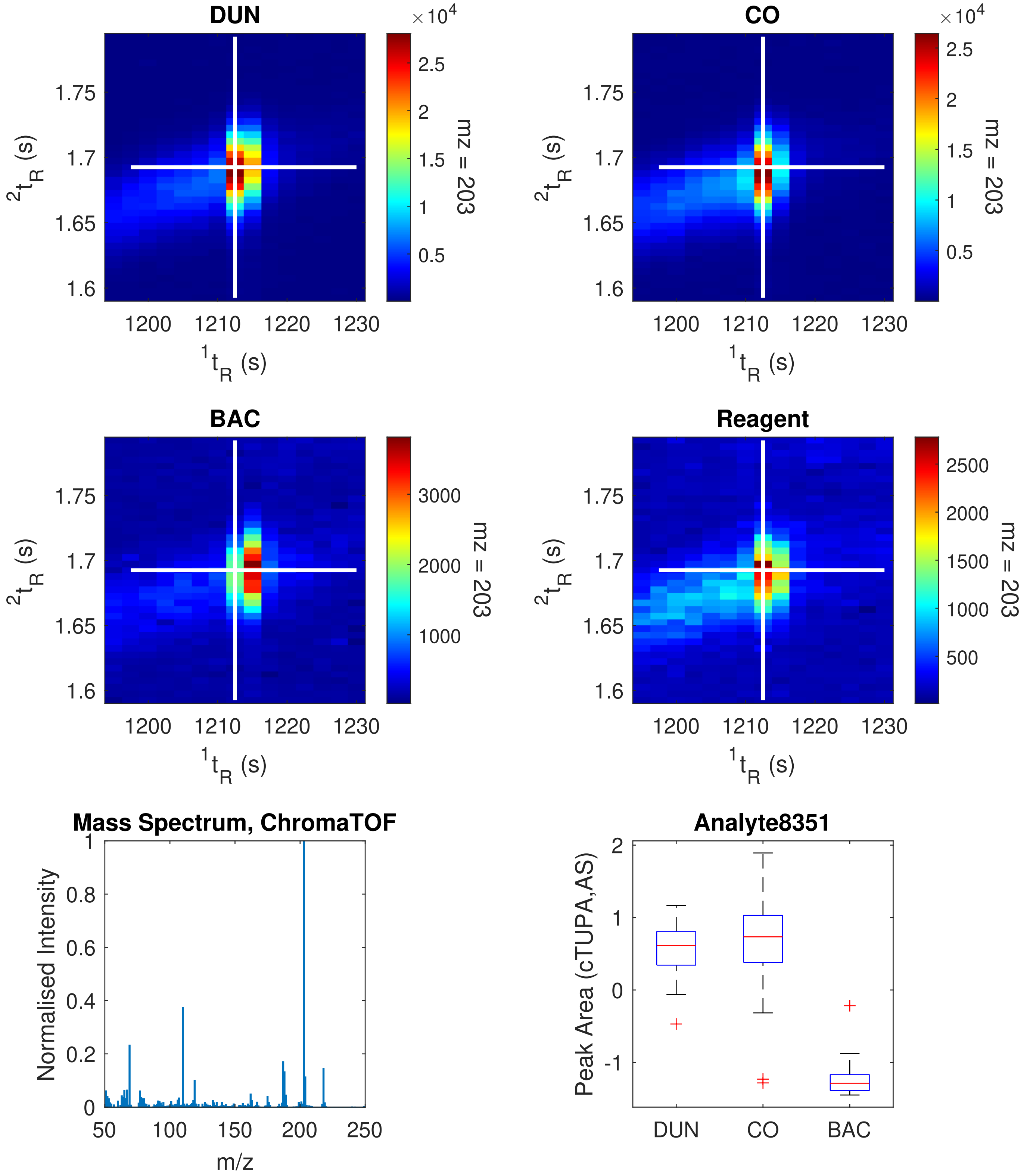

| Peak Information | Top Golm Metabolome Database Search Result |
| --- | --- |
| ${}^{1}t_R$(s): 1642.5 | |
| ${}^{2}t_R$(s): 1.905 | |
| Quantification Ion (m/z): 137 | 1-dotprod (Spectral Dissimilarity): Unknown |
| Analyte Name (ChromaTOF): Analyte13706 | Analyte Name (Library): Unknown |
| Retention Index (Observed): 1480.80 | Retention Index Difference (Library): Unknown |

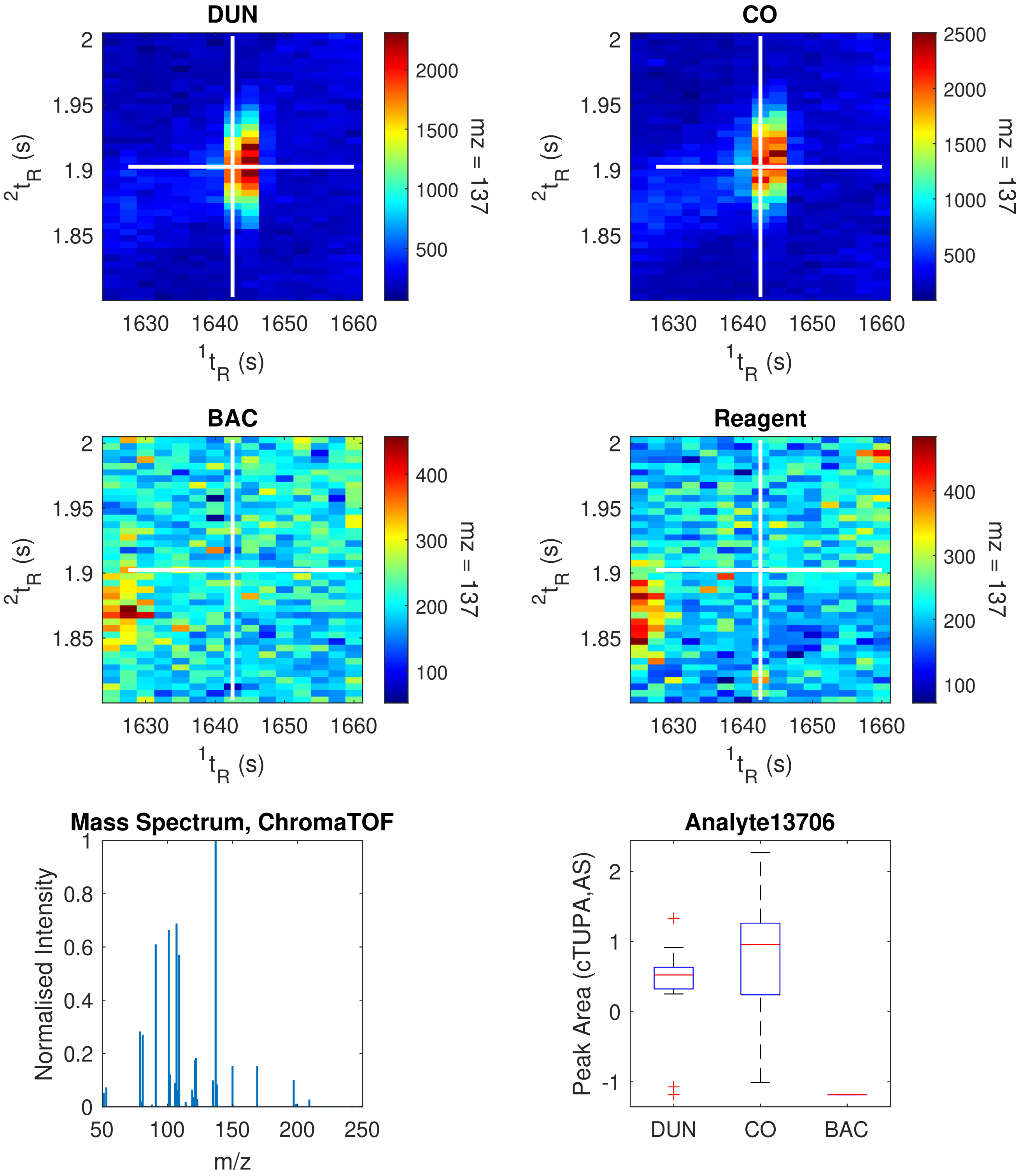

| Peak Information | Top Golm Metabolome Database Search Result |
|---|---|
| ${}^{1}t_R$(s): 1677.5 | |
| ${}^{2}t_R$(s): 1.270 | |
| Quantification Ion (m/z): 57 | 1-dotprod (Spectral Dissimilarity): 0.0846 |
| Analyte Name (ChromaTOF): Hexadecane | Analyte Name (Library): Pentadecane, n- |
| Retention Index (Observed): 1497.08 | Retention Index Difference (Library): 2.92 |

**Link:** http://gmd.mpimp-golm.mpg.de/Spectrums/e6650dda-783d-483f-bb3d-1d5c083da4ec.aspx
**Contributor:** Jäger C, Schomburg D Department of Bioinformatics and Biochemistry, Technische Universität Carolo-Wilhelmina Braunschweig, Langer Kamp 19B, D-38106 Braunschweig, Germany

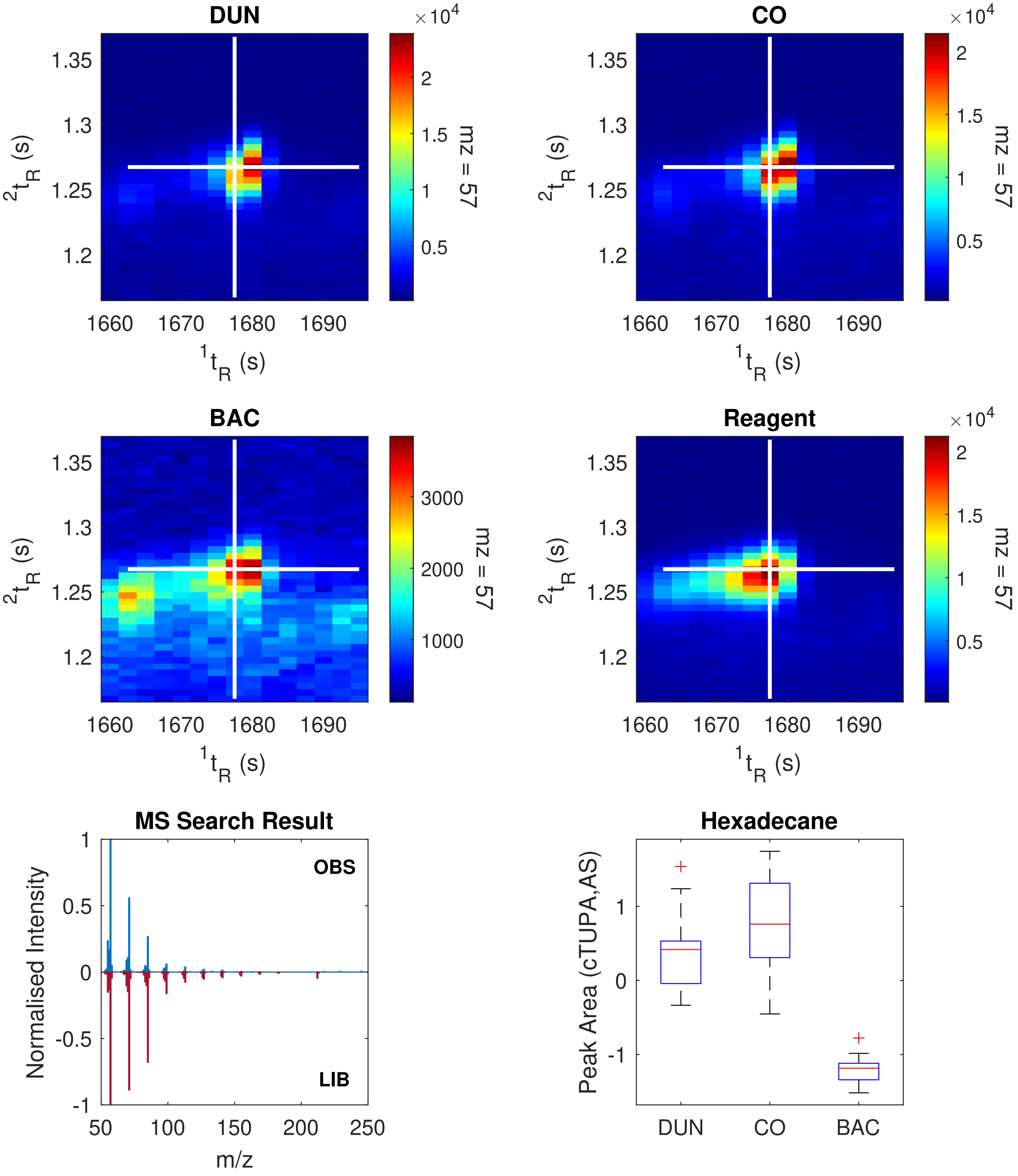

| Peak Information | Top Golm Metabolome Database Search Result |
|---|---|
| $^1t_R$(s): 1932.5 | |
| $^2t_R$(s): 1.480 | |
| Quantification Ion (m/z): 117 | 1-dotprod (Spectral Dissimilarity): 0.2868 |
| Analyte Name (ChromaTOF): Analyte16951 | Analyte Name (Library): 2-Aminoadipic-acid (2TMS) |
| Retention Index (Observed): 1624.02 | Retention Index Difference (Library): 3.79 |

**Link:** http://gmd.mpimp-golm.mpg.de/Spectrums/bec90e9c-eafd-4bec-9ad6-d2cde2ecdbbe.aspx
**Contributor:** Jäger C, Schomburg D Department of Bioinformatics and Biochemistry, Technische Universität Carolo-Wilhelmina Braunschweig, Langer Kamp 19B, D-38106 Braunschweig, Germany

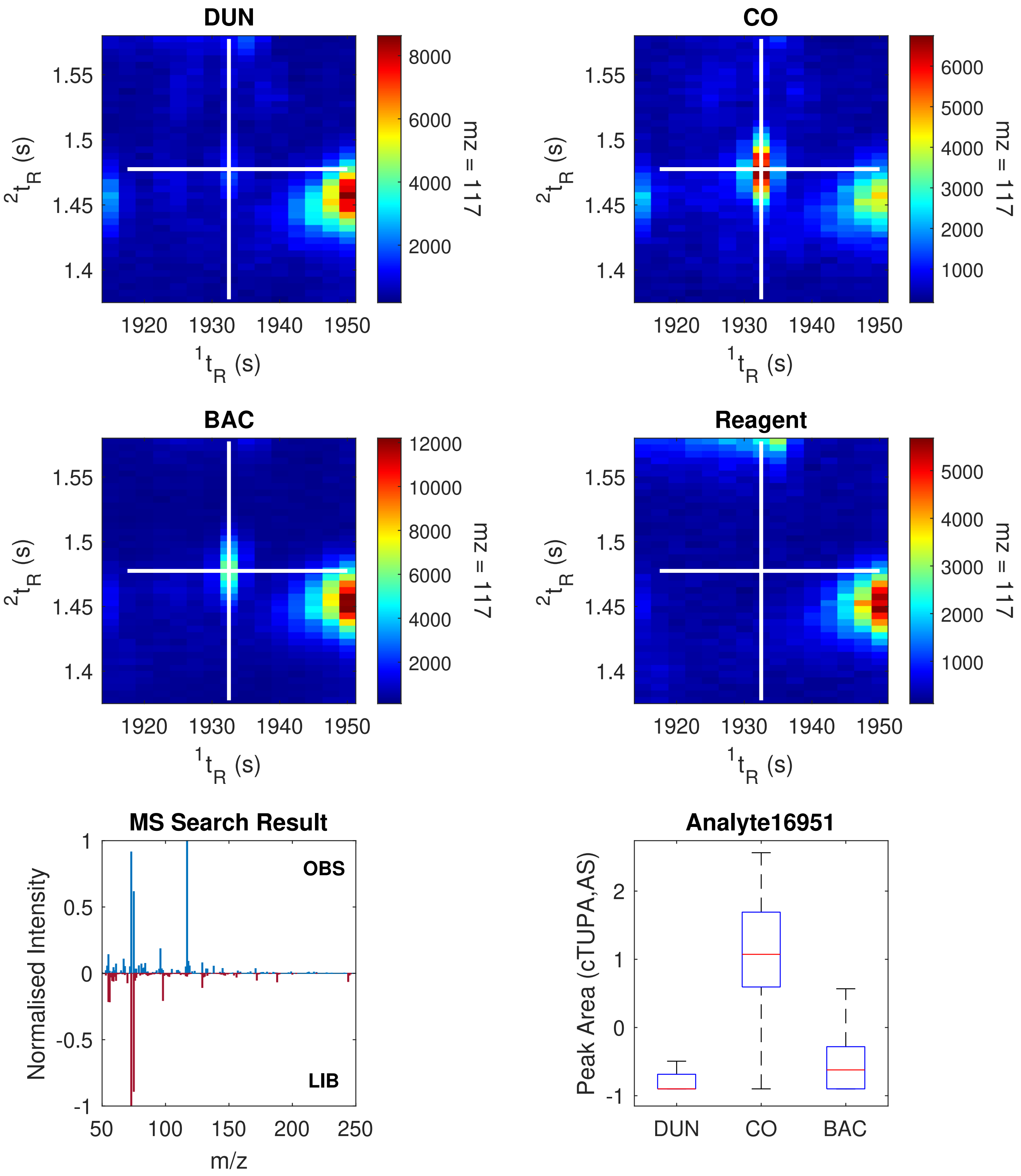

| Peak Information | Top Golm Metabolome Database Search Result |
|---|---|
| $^{1}t_R$(s): 2025.0 | |
| $^{2}t_R$(s): 1.990 | |
| Quantification Ion (m/z): 228 | 1-dotprod (Spectral Dissimilarity): 0.2952 |
| Analyte Name (ChromaTOF): Analyte17909 | Analyte Name (Library): Oxaloacetate (1MEOX) (3TMS) MP |
| Retention Index (Observed): 1672.07 | Retention Index Difference (Library): 0.64 |

**Link:** http://gmd.mpimp-golm.mpg.de/Spectrums/c0839970-b4b6-4089-9639-50f2aa8fd7d4.aspx
**Contributor:** Boelling C, Liebig F, Erban A, Kopka J, Max Planck Institute of Molecular Plant Physiology, Department of Molecular Plant Physiology (Prof. Willmitzer L), Am Muehlenberg 1, D-14476 Golm, Germany

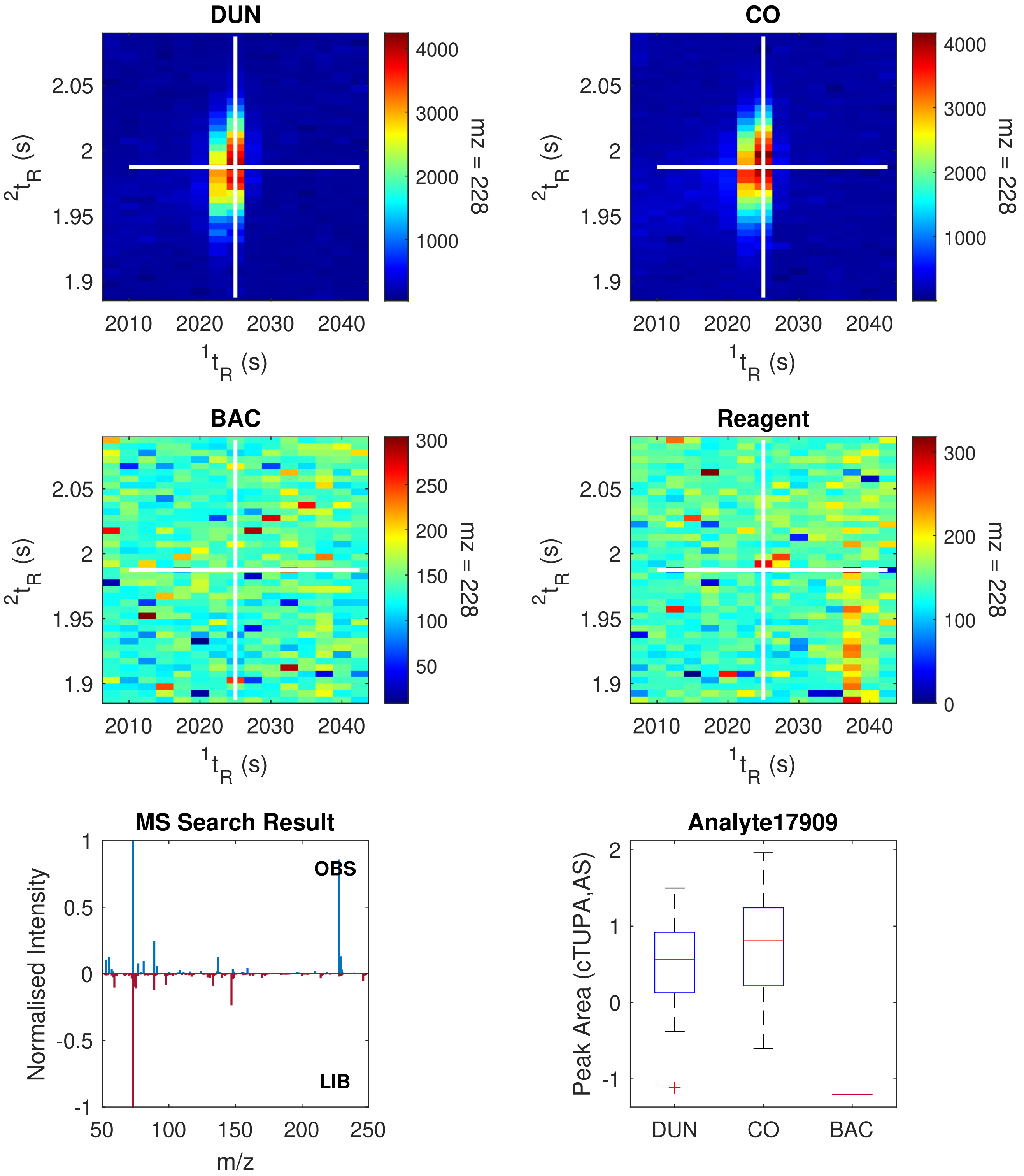

| Peak Information | Top Golm Metabolome Database Search Result |
|---|---|
| $^{1}t_R$(s): 2180.0 | |
| $^{2}t_R$(s): 1.600 | |
| Quantification Ion (m/z): 69 | 1-dotprod (Spectral Dissimilarity): Unknown |
| Analyte Name (ChromaTOF): Analyte19404 | Analyte Name (Library): Unknown |
| Retention Index (Observed): 1754.72 | Retention Index Difference (Library): Unknown |

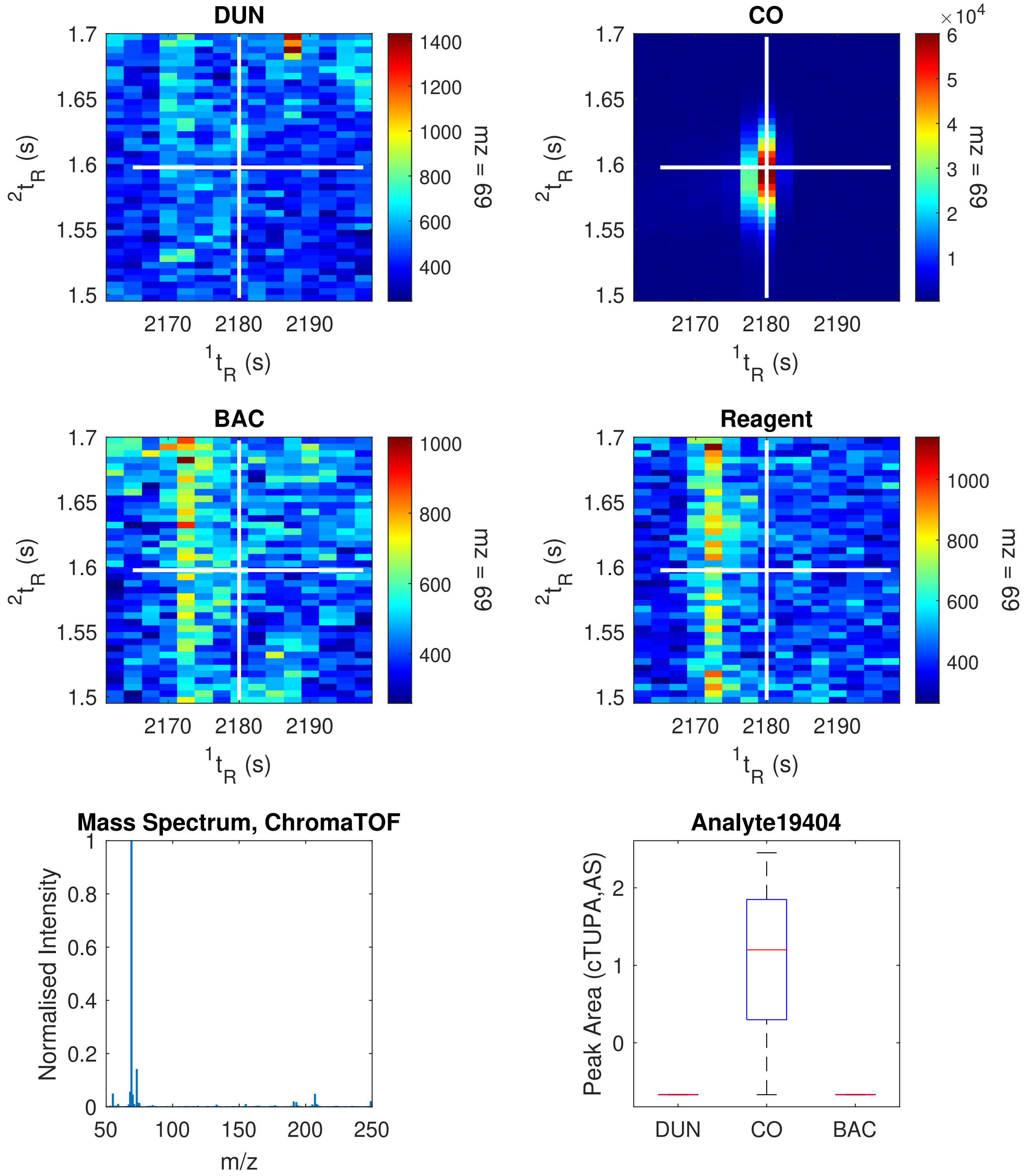

| Peak Information | Top Golm Metabolome Database Search Result |
|---|---|
| $^{1}t_R$(s): 2370.0 | |
| $^{2}t_R$(s): 1.365 | |
| Quantification Ion (m/z): 195 | 1-dotprod (Spectral Dissimilarity): 0.1310 |
| Analyte Name (ChromaTOF): Analyte21239 | Analyte Name (Library): Homoserine lactone, N-2-oxocaproyl- (1MEOX) (1TMS) BP |
| Retention Index (Observed): 1860.70 | Retention Index Difference (Library): 0.51 |

**Link:** http://gmd.mpimp-golm.mpg.de/Spectrums/7d96c6e2-42fc-46c8-85c2-0e11137c7210.aspx
**Contributor:** Boelling C, Liebig F, Erban A, Kopka J, Max Planck Institute of Molecular Plant Physiology, Department of Molecular Plant Physiology (Prof. Willmitzer L), Am Muehlenberg 1, D-14476 Golm, Germany

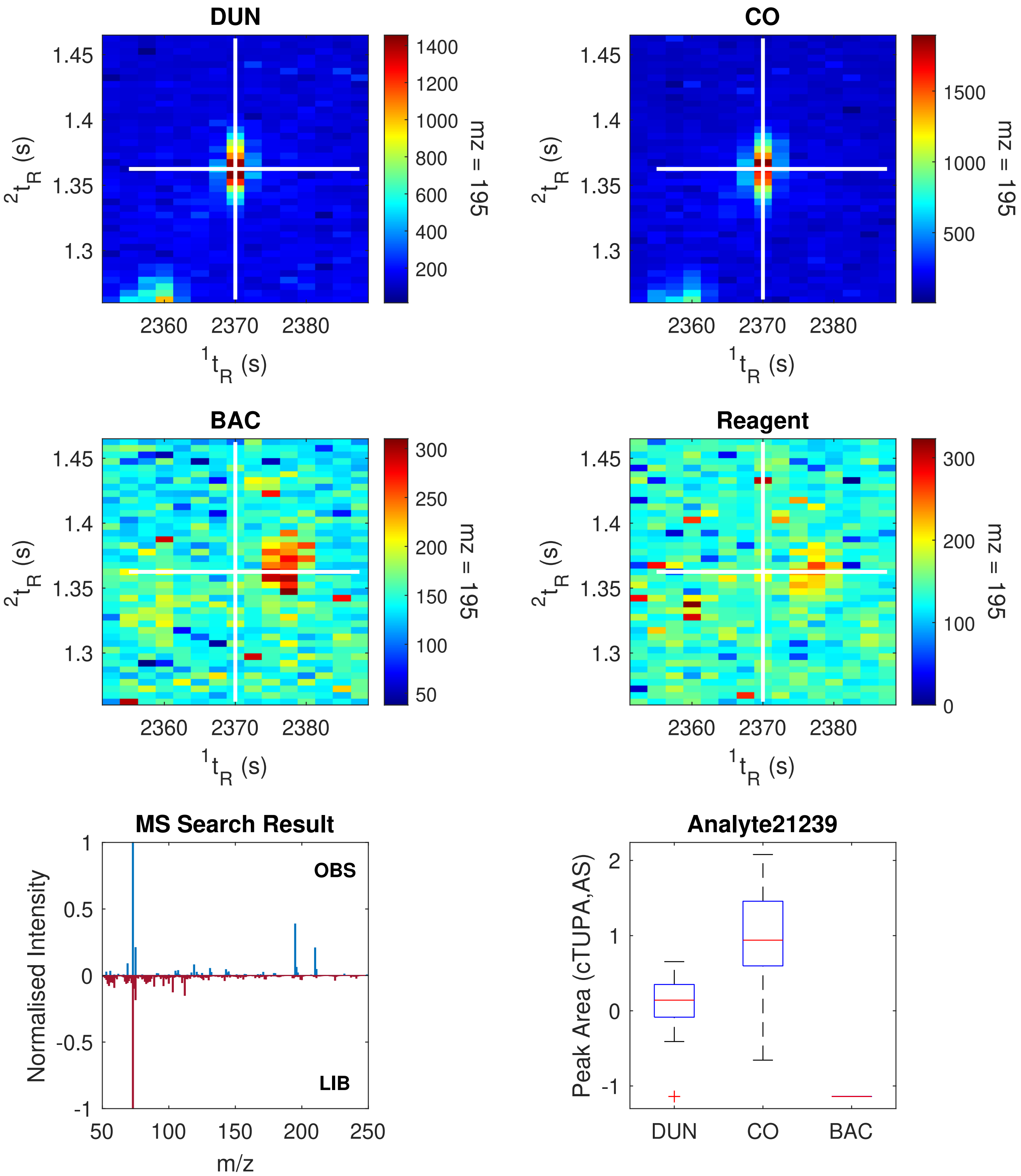

| Peak Information | Top Golm Metabolome Database Search Result |
|---|---|
| $^1t_R$(s): 2380.0 | |
| $^2t_R$(s): 1.775 | |
| Quantification Ion (m/z): 158 | 1-dotprod (Spectral Dissimilarity): Unknown |
| Analyte Name (ChromaTOF): Analyte21318 | Analyte Name (Library): Unknown |
| Retention Index (Observed): 1866.42 | Retention Index Difference (Library): Unknown |

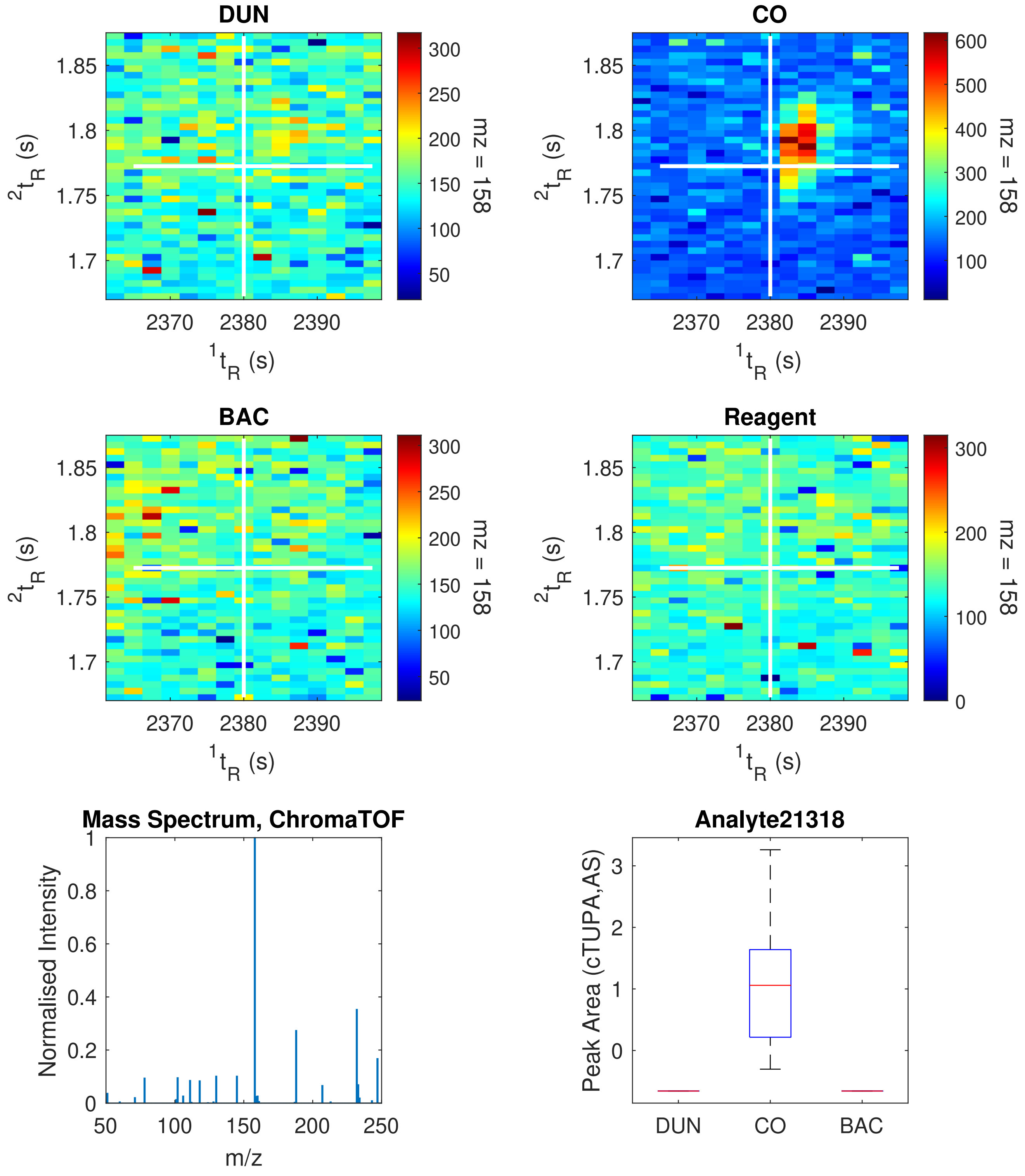

| Peak Information | Top Golm Metabolome Database Search Result |
|---|---|
| ${}^{1}t_R$(s): 2497.5 | |
| ${}^{2}t_R$(s): 1.320 | |
| Quantification Ion (m/z): 143 | 1-dotprod (Spectral Dissimilarity): 0.3436 |
| Analyte Name (ChromaTOF): Analyte22226 | Analyte Name (Library): Indole-3-acetic acid, 1H- (1TMS) |
| Retention Index (Observed): 1935.06 | Retention Index Difference (Library): 0.35 |

**Link:** http://gmd.mpimp-golm.mpg.de/Spectrums/35e55241-8b1a-42c3-a602-c1f05b31f818.aspx
**Contributor:** Liebig F, Erban A, Max Planck Institute of Molecular Plant Physiology, Department of Molecular Plant Physiology (Prof. Willmitzer L), Am Muehlenberg 1, D-14476 Golm, Germany

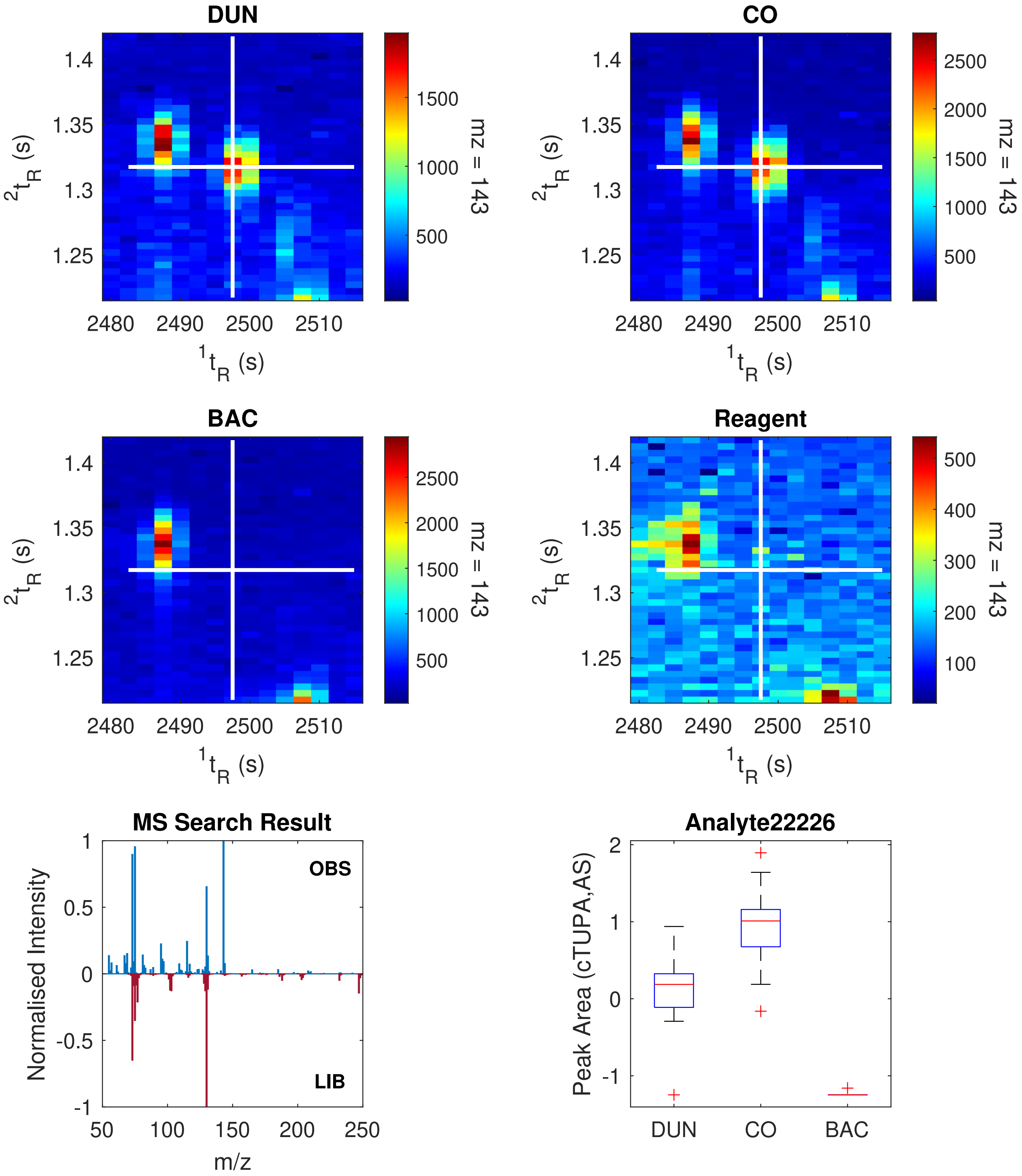

| Peak Information | Top Golm Metabolome Database Search Result |
|---|---|
| $^1t_R$(s): 2597.5 | |
| $^2t_R$(s): 1.335 | |
| Quantification Ion (m/z): 165 | 1-dotprod (Spectral Dissimilarity): 0.2324 |
| Analyte Name (ChromaTOF): Analyte23041 | Analyte Name (Library): Pyruvic acid, 4-hydroxyphenyl- (1MEOX) (3TMS) MP |
| Retention Index (Observed): 1994.76 | Retention Index Difference (Library): 1.63 |

**Link:** http://gmd.mpimp-golm.mpg.de/Spectrums/14c81dc4-1c9f-4731-a2cd-974fdd692d47.aspx
**Contributor:** Boelling C, Liebig F, Erban A, Kopka J, Max Planck Institute of Molecular Plant Physiology, Department of Molecular Plant Physiology (Prof. Willmitzer L), Am Muehlenberg 1, D-14476 Golm, Germany

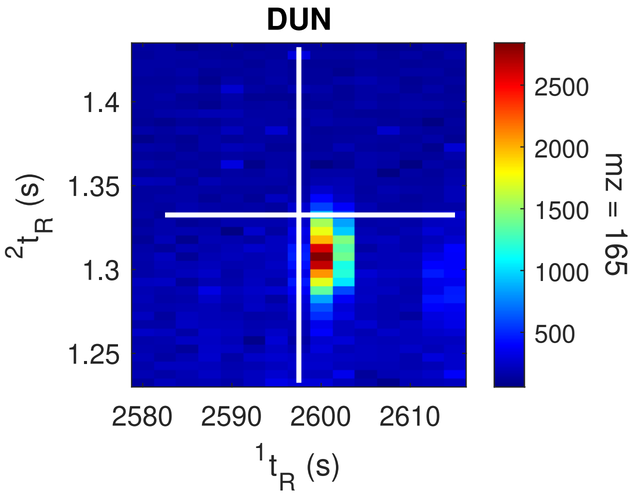


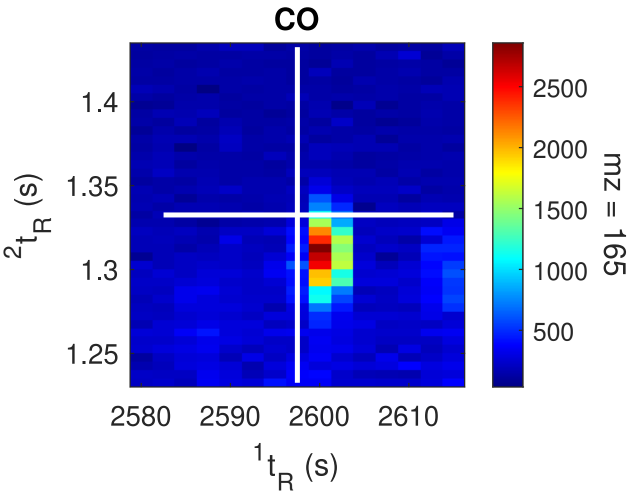


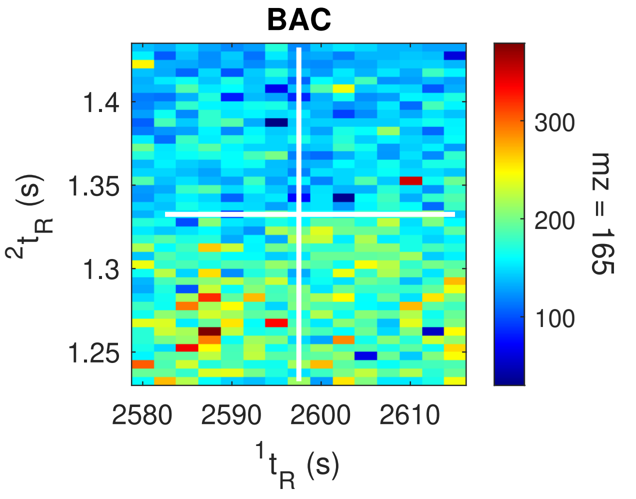


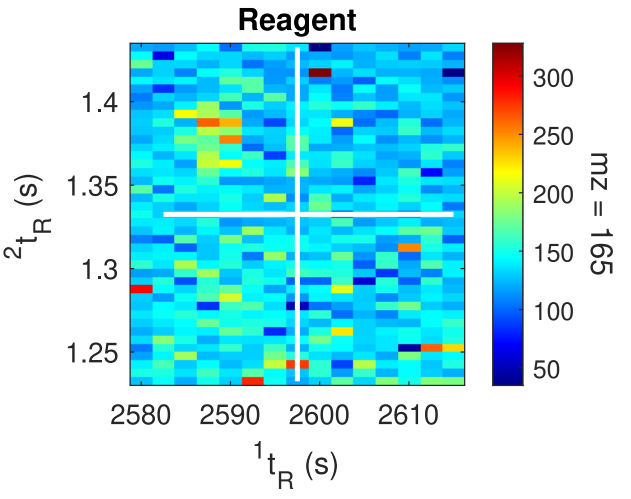


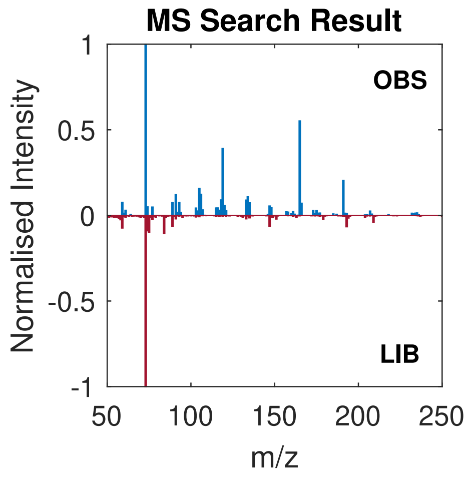


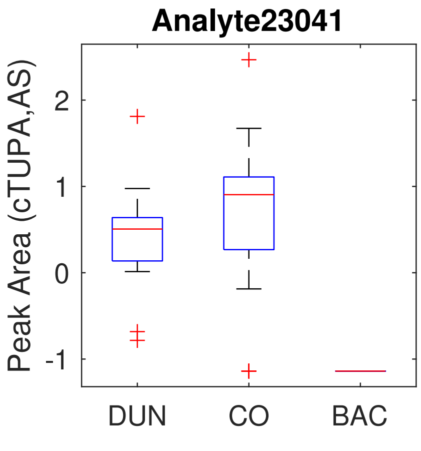

| Peak Information | Top Golm Metabolome Database Search Result |
|---|---|
| $^1t_R$(s): 2645.0 | |
| $^2t_R$(s): 2.045 | |
| Quantification Ion (m/z): 376 | 1-dotprod (Spectral Dissimilarity): 0.0793 |
| Analyte Name (ChromaTOF): Analyte23397 | Analyte Name (Library): Gluconic acid, 2-amino-2-deoxy- (7TMS) |
| Retention Index (Observed): 2024.59 | Retention Index Difference (Library): 1.14 |

**Link:** http://gmd.mpimp-golm.mpg.de/Spectrums/bfddda38-5af0-4dc1-a200-152b9380b8d4.aspx
**Contributor:** Boelling C, Liebig F, Erban A, Kopka J, Max Planck Institute of Molecular Plant Physiology, Department of Molecular Plant Physiology (Prof. Willmitzer L), Am Muehlenberg 1, D-14476 Golm, Germany

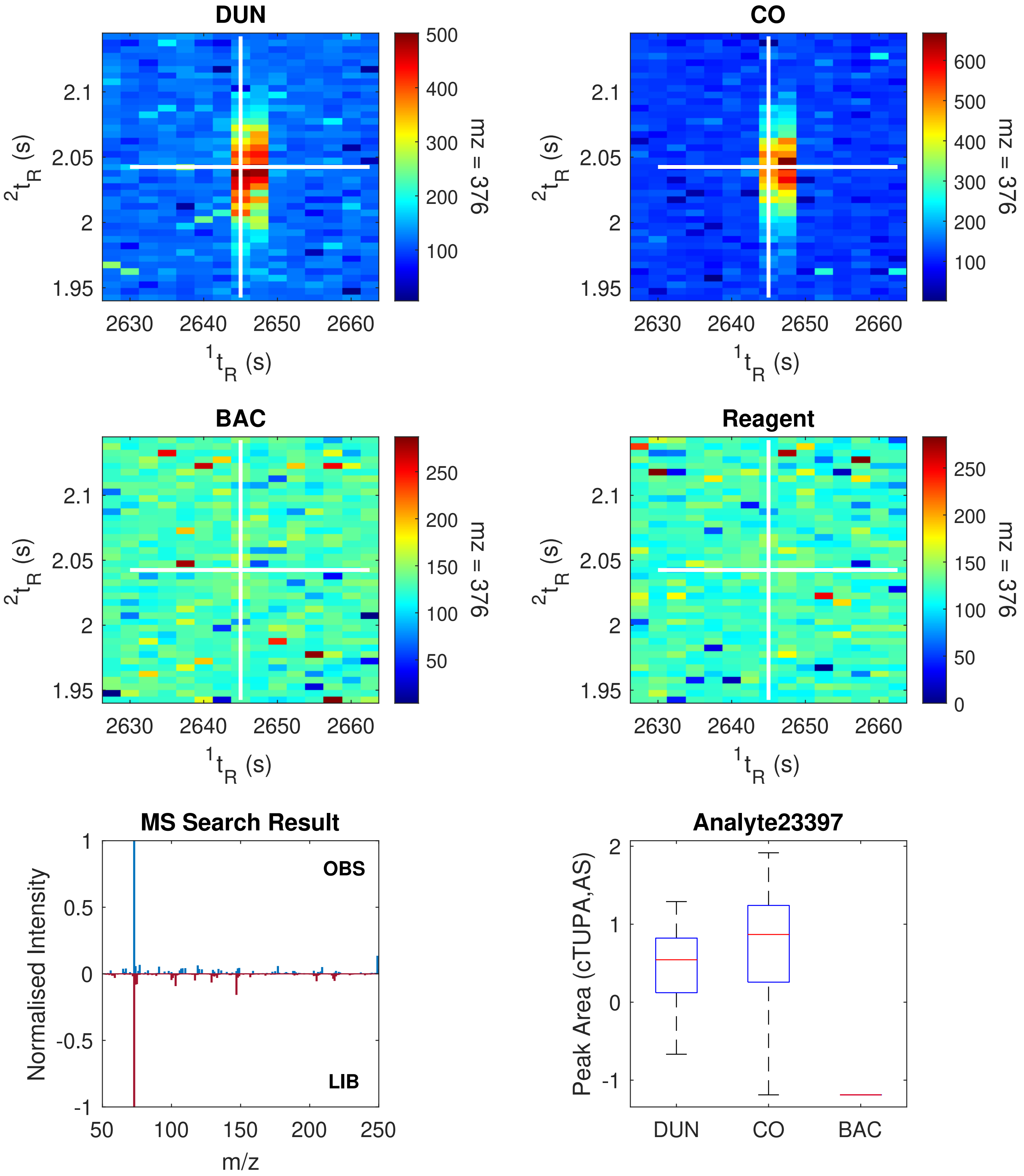

| Peak Information | Top Golm Metabolome Database Search Result |
|---|---|
| $^{1}t_R$(s): 2832.5 | |
| $^{2}t_R$(s): 1.865 | |
| Quantification Ion (m/z): 80 | 1-dotprod (Spectral Dissimilarity): Unknown |
| Analyte Name (ChromaTOF): Analyte24829 | Analyte Name (Library): Unknown |
| Retention Index (Observed): 2145.06 | Retention Index Difference (Library): Unknown |

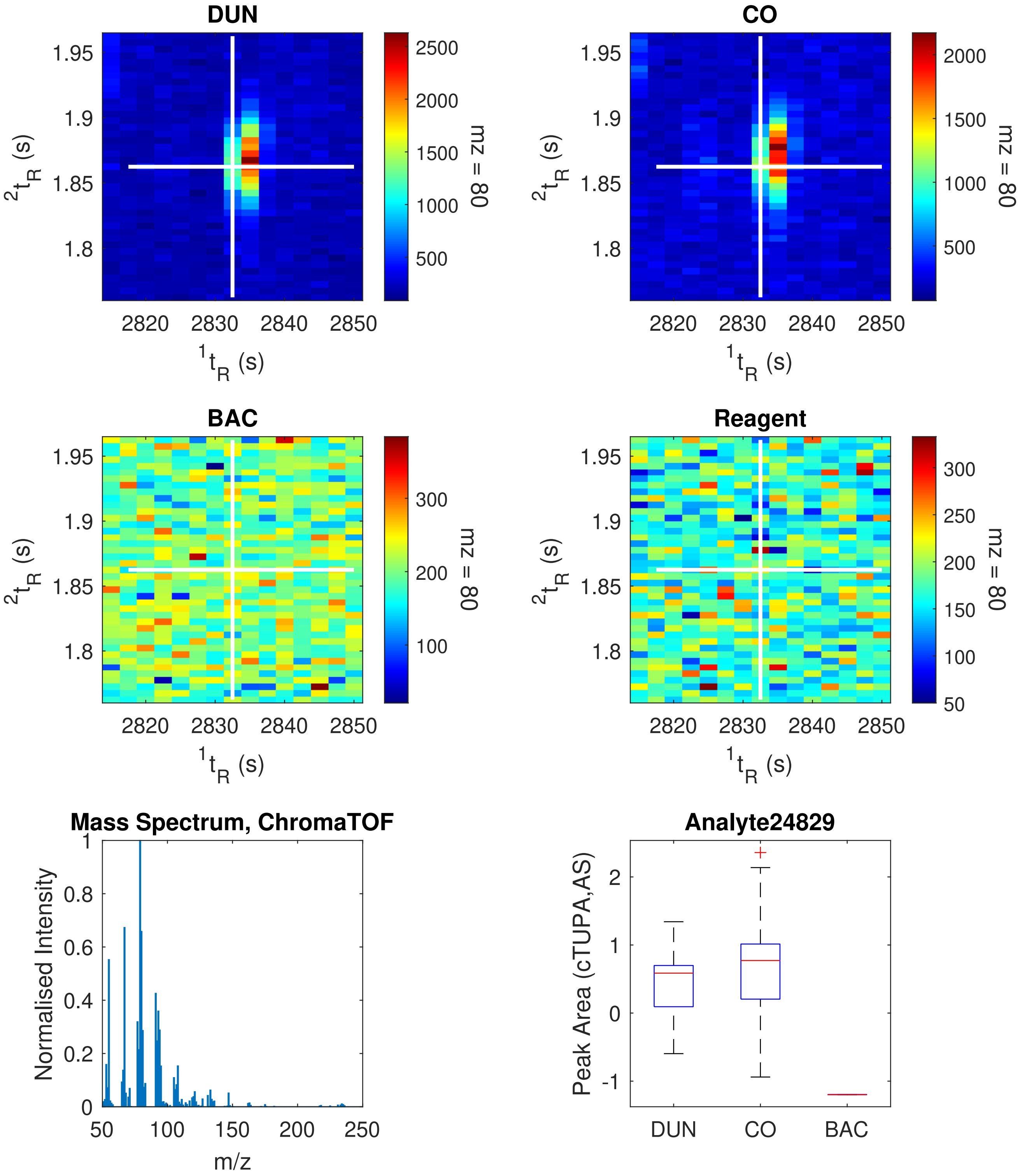

| Peak Information | Top Golm Metabolome Database Search Result |
|---|---|
| $^1t_R$(s): 3200.0 | |
| $^2t_R$(s): 1.415 | |
| Quantification Ion (m/z): 233 | 1-dotprod (Spectral Dissimilarity): 0.0553 |
| Analyte Name (ChromaTOF): Analyte27584 | Analyte Name (Library): NA |
| Retention Index (Observed): 2334.57 | Retention Index Difference (Library): 1.87 |

**Link:** http://gmd.mpimp-golm.mpg.de/Spectrums/3ffe8941-e355-4f09-a740-ec684f396a2f.aspx
**Contributor:** Kopka J, Max Planck Institute of Molecular Plant Physiology, Department of Molecular Plant Physiology (Prof. Willmitzer L), Am Muehlenberg 1, D-14476 Golm, Germany

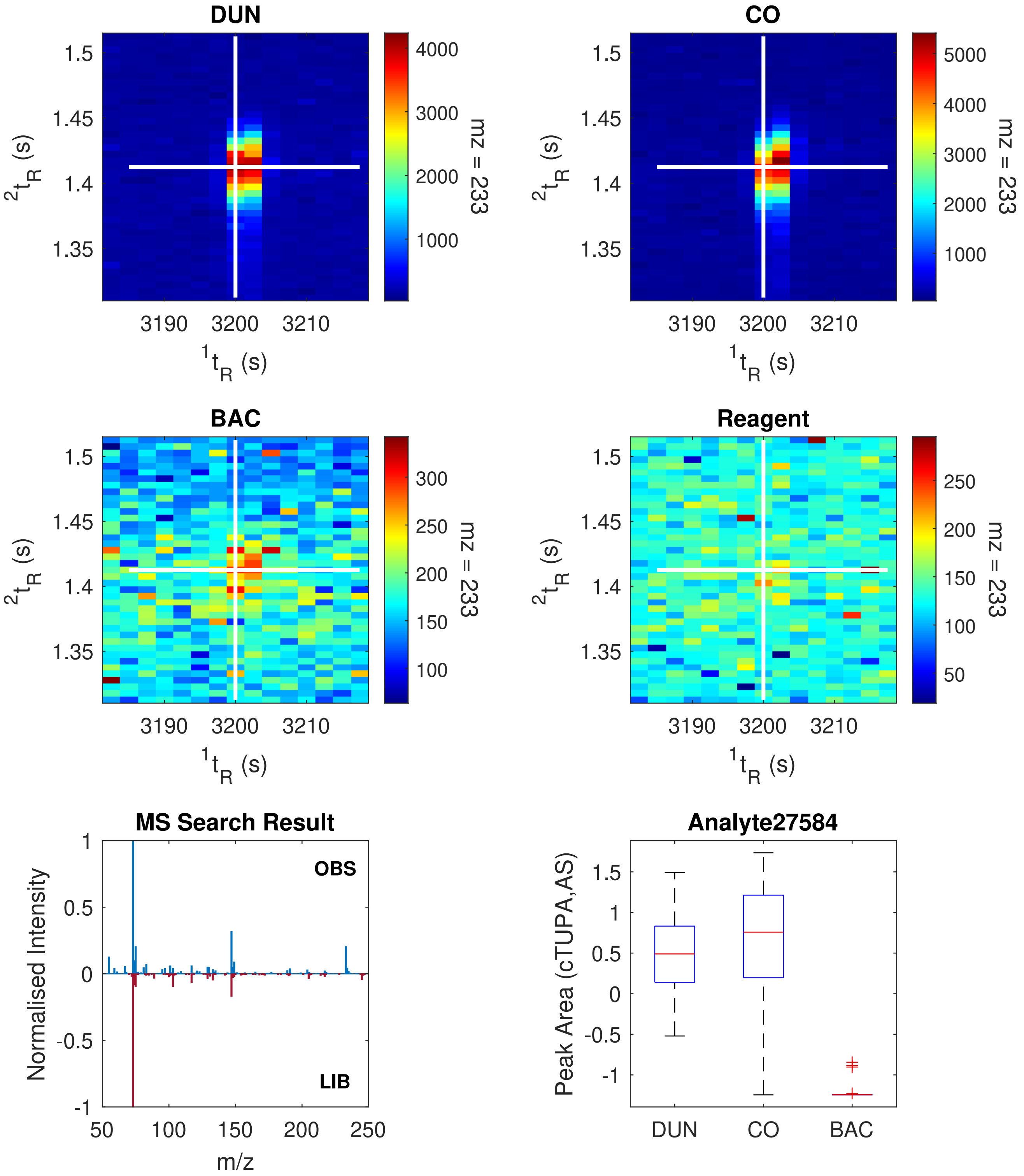

| Peak Information | Top Golm Metabolome Database Search Result |
|---|---|
| ${}^{1}t_R$(s): 3242.5 | |
| ${}^{2}t_R$(s): 1.475 | |
| Quantification Ion (m/z): 195 | 1-dotprod (Spectral Dissimilarity): 0.3819 |
| Analyte Name (ChromaTOF): Analyte27833 | Analyte Name (Library): Galactose-6-phosphate (1MEOX) (6TMS) BP |
| Retention Index (Observed): 2416.04 | Retention Index Difference (Library): 0.42 |

**Link:** http://gmd.mpimp-golm.mpg.de/Spectrums/84406ec0-9f61-42c0-858c-37079019ec05.aspx
**Contributor:** Erban A, Strehmel N, Kopka J, Max Planck Institute of Molecular Plant Physiology, Department of Molecular Plant Physiology (Prof. Willmitzer L), Am Muehlenberg 1, D-14476 Golm, Germany

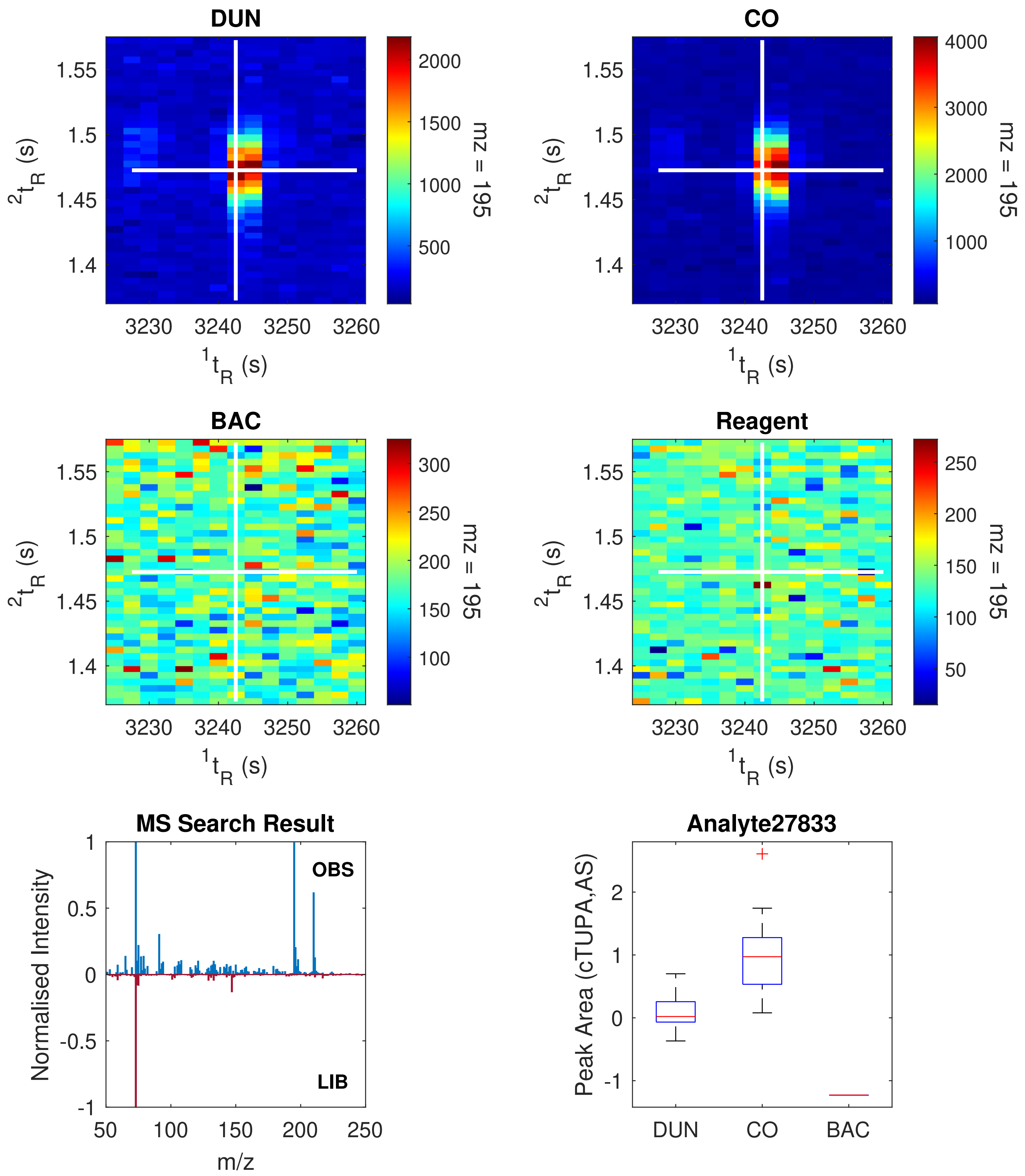

| Peak Information | Top Golm Metabolome Database Search Result |
|---|---|
| $^{1}t_R$(s): 4215.0 | |
| $^{2}t_R$(s): 1.840 | |
| Quantification Ion (m/z): 324 | 1-dotprod (Spectral Dissimilarity): 0.3138 |
| Analyte Name (ChromaTOF): Analyte33374 | Analyte Name (Library): Cycloeucalenol (1TMS) |
| Retention Index (Observed): Undefined | Retention Index (Library): Undefined |

**Link:** http://gmd.mpimp-golm.mpg.de/Spectrums/e68eba0e-3d8c-44a8-8bbe-0001a2b2c23f.aspx
**Contributor:** Moritz T, Umea Plant Science Centre, Department of Forest Genetics and Plant Physiology, Swedish University of Agricultural Sciences, SE-901 83 Umea, Sweden

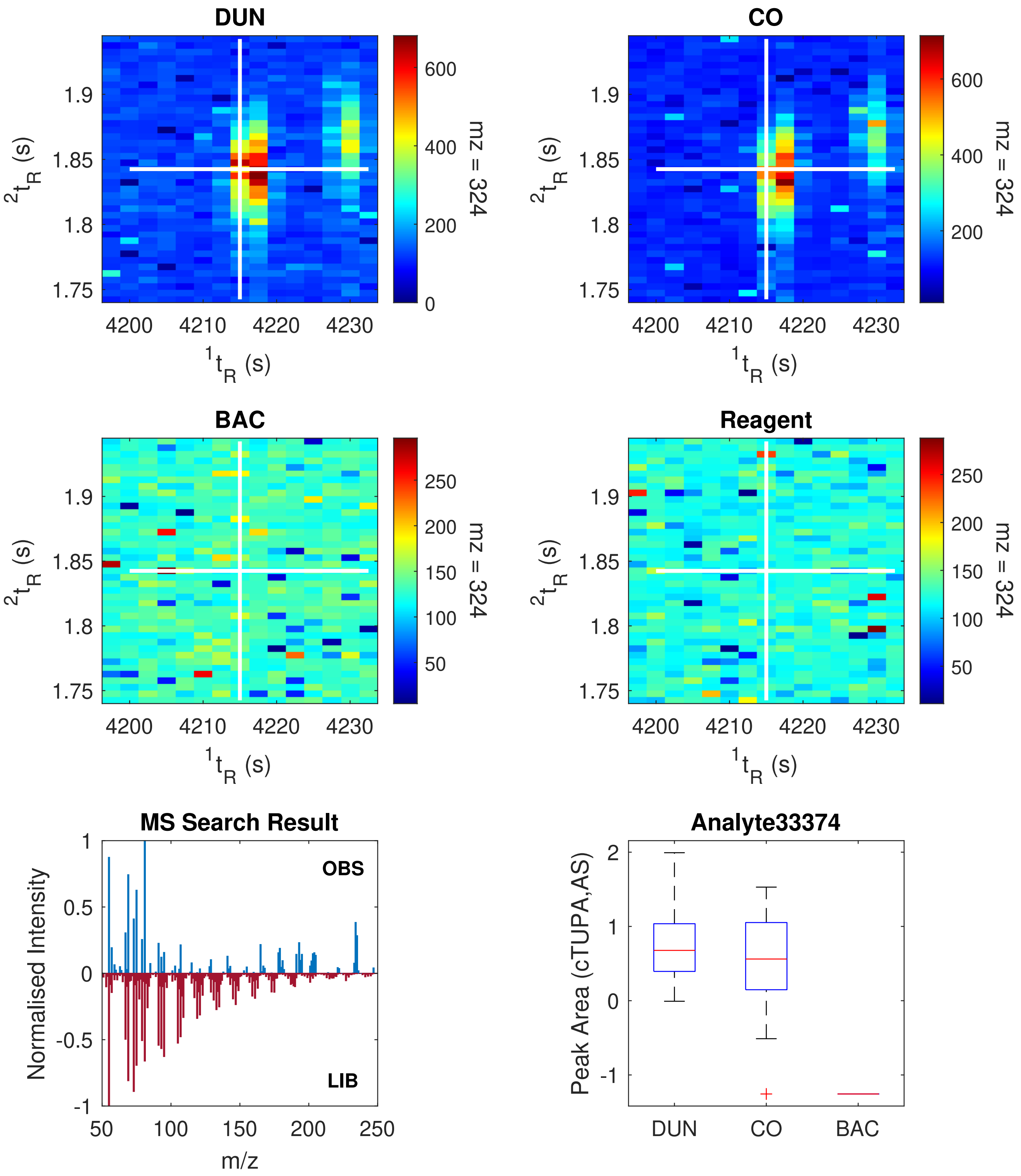
